# VTA GABA Cell Bipotential Induction of iLTP or iLTD Synaptic Plasticity is Input Selective, where iLTD is Uniquely Eliminated by Cocaine

**DOI:** 10.1101/2025.10.06.680313

**Authors:** Bridget J. Wu, Seth Hoffman, Hannah Urey, Isaac Forbush, Devin Bird, Craig Brailsford, Jeffrey G. Edwards

## Abstract

The ventral tegmental area (VTA) is a key reward circuit hub, implicated in drug seeking and addictive behaviors. The VTA contains dopaminergic and GABAergic neurons that both play roles in reward prediction, aversion, motivated reward behavior, etc. Synaptic plasticity, including VTA excitatory and inhibitory long-term potentiation (LTP/iLTP) and long-term depression (LTD/iLTD), are fundamentally involved in processing reward learning and memory, which is maladaptively altered by abused drugs mediating dependence induction. This report extends our prior research of the understudied VTA GABA cells and the rationale for their bipotential plasticity (iLTP or iLTD) capacity by optogenetic circuit level examination of their unique GABAergic inputs. In addition, we examine potential cocaine impact on both plasticity forms. Optogenetic activation of either lateral hypothalamus or rostromedial tegmental nucleus induced iLTP in VTA GABA cells, while optogenetic activation of local VTA GABAergic inputs induced iLTD. This suggests expression of bipotential plasticity is input specific, and highlights implications for each type of plasticity on reward signaling. Drug of abuse cocaine eliminated iLTD while sparing iLTP, suggesting selective impairment of local GABA signaling to VTA GABA cells by cocaine versus projection GABA signaling. This emphasizes potential differential VTA GABA cell plasticity in reward processing and the need to further examine cocaine impact on inhibitory signaling in addition to known impact on the dopaminergic system. By elucidating the circuit-dependence of plasticity type in VTA GABA cells and drug-induced impact, our research could potentially identify additional targets for therapeutic intervention of drug dependence via the GABAergic system.

## Introduction

As an essential component of the mesolimbic dopamine system, the ventral tegmental area (VTA) plays a pivotal role in reward, drug seeking, withdrawal and reinstatement (Koob and Volkow, 2010). VTA functionality is therefore central to the neurobiological understanding of addiction in substance use disorders, where normal reward processing is hijacked by exogenous substances, leading to maladaptive patterns of behavior and synaptic plasticity. Within this circuitry, dopamine (DA) neurons are affected by drug exposure, leading to drug-dependent states (Juarez and Han, 2016; Lüscher and Ungless, 2006; Ungless et al., 2001). Equally important, yet less studied, are the VTA GABAergic neurons, which are composed of both projection and interneurons subtypes. The VTA GABA interneurons innervate and modulate dopamine activity and release (Klitenick et al., 1992), impacting reward *in vivo* (Bocklisch et al., 2013; van Zessen et al., 2012), as well as roles in self-stimulation reward paradigms (Lassen et al., 2007), conditioned place aversion (Tan et al., 2012), conditioned place preference (Bocklisch *et al*., 2013) and associative reward learning (Brown et al., 2012). The projecting VTA GABA neurons function independently of direct action on DA cells and instead project to nucleus accumbens (NAc) where they enhance associative reward learning (Brown *et al*., 2012), and to the ventral pallidum where they encode reward value and motivation (Zhou et al., 2022), among other targets. These inhibitory interneurons also ensure a delicate balance between excitation and inhibition, modulating dopamine release in response to salient reward/aversive stimuli (Creed et al., 2014; Erhardt et al., 2002; Omelchenko et al., 2009; van Zessen *et al*., 2012). Disruptions to this balance, particularly through alterations in synaptic plasticity induced by drug exposure, contribute significantly to the development and perpetuation of addictive behaviors (Dacher and Nugent, 2011; Edwards et al., 2017; Lüscher and Malenka, 2011; Nugent et al., 2007).

Synaptic plasticity is a crucial process for the brain to adapt to new information, enabling the formation of new memories. There are two main types of plasticity, long-term potentiation (LTP), and long-term depression (LTD). LTP involves strengthening of a synapse while LTD weakens a synapse, which occurs at inhibitory GABAergic synapses as well (iLTP and iLTD). All these plasticity types can be impacted by exposure to drugs of abuse (Kasanetz et al., 2010; Shen and Kalivas, 2013). Drug-induced plasticity changes were studied extensively at synapses involving dopamine neurons. Less is known regarding drug impact on VTA GABA cells until Bocklisch et al. (2013) demonstrated an iLTP in VTA GABA cells from optogenetically activated NAc inputs using high frequency stimulus. This iLTP was presynaptic, PKA-dependent, and occluded by 5 days of cocaine administration. Our recent published study examining non-selective GABA inputs to VTA GABA cells demonstrated a 5-Hz stimulus induced either iLTP or iLTD in VTA GABA neurons (Nufer et al., 2023). While the iLTD was GABA_B_ receptor-dependent, the iLTP presented a more complex picture, being only partially contingent upon N-methyl-D-aspartic acid receptor (NMDAR) involvement (Nufer *et al*., 2023), and thus other receptor signaling could be involved.

A recent study reported that among VTA DA neurons somatodendritic release of cholecystokinin (CCK) underlies iLTP of specific GABAergic inputs, which was eliminated by CCK_2_ receptors antagonists (Martinez Damonte et al., 2023). This suggests CCK could also be involved in iLTP of VTA GABA cells. In addition to CCK, we revisited the nitric oxide synthase (NOS) pathway, which plays a crucial role in the initiation of iLTP at VTA GABA to DA synapses as well (Nugent *et al*., 2007). Our previous study illustrated SNAP, a NO donor, potentiated 50% of the VTA GABA neurons (Nufer *et al*., 2023). We therefore investigated NOS involvement in VTA GABA cell iLTP.

The bipotential plasticity phenomenon is rare with few notations in the brain including VTA DA neurons where GABAergic inputs from RMTg exhibit iLTD and periductal gray inputs exhibit iLTP (St Laurent and Kauer, 2019). VTA GABA neurons receive GABA inputs from various regions, including the NAc (Bocklisch *et al*., 2013; Edwards *et al*., 2017), lateral habenula (LHb), laterodorsal tegmental nucleus, etc. (Faget et al., 2016; Omelchenko *et al*., 2009), lateral hypothalamus (LH) Nieh, 2016 #3028} and rostromedial tegmental nucleus (RMTg) Barrot, 2012 #1873}(Jhou et al., 2009). We previously demonstrated iLTP and iLTD to VTA GABA cells is presynaptic (Nufer *et al*., 2023), leading us to hypothesize that these distinct plasticity forms could be attributable to the origin of inhibitory inputs to VTA GABA neurons, which we examined here using optogenetics. However, given the heterogeneity of VTA GABA cells (Phillips et al., 2022), a complementary hypothesis might lie in the existence of GABAergic neuron subtypes identified within the VTA (Margolis et al., 2012; Morales and Margolis, 2017; Paul et al., 2019; Phillips *et al*., 2022; Root et al., 2020), each uniquely predisposed to different plasticity responses. We investigated this hypothesis by studying the distribution of the GABA synthesis enzymes glutamic acid decarboxylase 67-kD (GAD 67) and glutamic acid decarboxylase 65-kD (GAD65) in VTA GABA neurons and their displayed plasticity type.

Lastly, as drug-induced modification to endogenous plasticity is correlated to drug dependence induction in DA neurons (Lüscher and Malenka, 2011), it is essential to investigate drugs of abuse impact on the VTA GABAergic circuit to provide a more complete picture of drug impact on the reward circuit, including the inhibitory circuit. This includes cocaine, which alters endogenous DA cell plasticity (Luscher and Bellone, 2008; Ungless *et al*., 2001) and iLTP of GABA inputs to VTA GABA cells (Bocklisch *et al*., 2013). However, cocaine impact on iLTD is still unknown. Therefore, we also examined acute and chronic impact on VTA GABA cell plasticity with a focus on iLTD as it is GABA_B_ receptor-dependent plasticity (Nufer *et al*., 2023) as cocaine impacts GABA_B_ function including cue-induced drug seeking and cocaine sensitization (Di Ciano and Everitt, 2003; Jayaram and Steketee, 2004; Yamaguchi et al., 2002).

## Methods

All experiments were performed in accordance with Institutional Animal Care and Use Committee protocols, following National Institutes of Health Guide for the care and use of laboratory animals. Experiments were approved by the Brigham Young University Institutional Animal Care and Use Committee, Animal Welfare Assurance Number A3783-01.

### Animals

GAD67/GFP^+^ mice were bred inhouse for >10 years (Jackson Laboratory, stock number: 007677, strain code: CB6-Tg(Gad1-EGFP)G42Zjh/J). Other mice purchased from Jackson Laboratory include VGAT::IRES-Cre (Jackson Laboratory, stock number: 028862, strain code: B6J.129S6(FVB)-*Slc32a1^tm2(Cre)Lowl^*) and GAD65 mCherry^+^ (Jackson Laboratory, stock number: 023140, Strain code: B6;129S-*Gad2^tm1.1Ksvo^*/J). VGAT::IRES-Cre/GAD67 GFP^+^ and GAD67 GFP^+^/GAD65 mCherry^+^ lines were bred in house. A VGAT::IRES-Cre/GAD67 GFP^+^ line was established by crossing VGAT::IRES-Cre males with GAD67 GFP^+^ females. A GAD67 GFP^+^/GAD65 mCherry^+^ line was produced by crossing GAD65 mCherry^+^ males with GAD67 GFP^+^ females. Male and female GAD67 GFP^+^ used in CCK and SNAP experiments as well as male and female GAD67 GFP^+^/GAD65 mCherry^+^ were juvenile mice between postnatal 15 to 30 days. Since post-surgery recovery after stereotaxis injection would normally take 3 to 8 weeks, animals for all the optogenetic experiments were young adults between postnatal 60 to 90 days. Cocaine-injected mice were used at postnatal 20-30 days, except those used for the optogenetic experiments, which again needed recovery times and thus age-matched to other optogenetic experiments between postnatal days 60 to 90. Mice were maintained on a 12-hour light/dark cycle and provided with food and water *ad libitum*.

### Stereotaxic injections

VGAT::IRES-Cre/GAD67 GFP^+^ mice (male and female) between prenatal day 35-45 received stereotaxic microinjection in a Kopf stereotaxic apparatus (Model 940). Mice were anesthetized using 1-3% isoflurane dispensed with a Kent SomnoSuite. Carprofen tablets were administered 24 hours before and for at least 72 hours following surgery, as well as a subcutaneous injection of buprenorphine (0.1 mg/kg) after induction of anesthesia and immediately before surgery. AAV2/1-EF1a-DIO-hChR2(H134R)-mCherry (UNC Vector Core, North Carolina) was bilaterally injected into the Lateral Hypothalamus (AP –1.3, ML ±0.6 to 0.8, DV –5.2 to –5.3; 1 µL), RMTg (AP –3.5, ML ±0.5, DV –4.4; 500 nL) and VTA (AP –2.5, ML ±0.5, DV –4.4; 500 nL), respectively. Animal recovery and viral incubation lasted 3 to 8 weeks post-surgery, after which time transverse slices containing the VTA were prepared for electrophysiology or immunohistochemistry.

### Immunohistochemistry (IHC)

Immunohistochemistry confirmed viral injection location by confirming ChR2-mCherry expression. Mice used for IHC were male/female post-surgery VGAT::IRES-Cre/GAD67 GFP^+^ (P60-90) mice. Brains were transcardially perfused with 0.1 M phosphate-buffered saline (PBS) followed by 4% paraformaldehyde in 0.1 M PBS (pH 7.4). Brains were dissected out, cryoprotected in 15% sucrose solution for 30 minutes, 20% sucrose solution for 1 hour, and 30% sucrose solution for 24 hours, frozen to –80 °C and sliced into 15-30 μm sections, then collected into 0.1 M PBS using a free-floating staining procedure. For examination of tyrosine hydroxylase (TH), slices were permeabilized with 0.2% Triton-X (Fisher Bioreagents) for 30 minutes, washed with 5% normal goat serum and 1% bovine serum albumin in 0.1 M PBS for 2 hours, and treated with primary antibody for anti-tyrosine hydroxylase (TH) (1:1000; sheep polyclonal; AB2492277; Phosphosolutions) in 5% normal goat serum and 1% bovine serum albumin in PBS overnight at 4°C. Slices were then washed twice with 0.1 M PBS, followed by one wash of 0.2% Triton-X (Fisher Bioreagents) in 0.1 M PBS for 30 minutes, and one wash of 5% normal goat serum and 1% bovine serum albumin in 0.1 M PBS for 2 hours. After treatment of anti-sheep secondary antibody (1:500, AlexaFluor 405) in 5% normal goat serum and 1% bovine serum albumin in PBS for 2 hours at room temperature, slices were washed three times with tris-buffered saline and mounted onto Superfrost Plus microscope slides (VWR). After drying overnight, slides were cover slipped with DAPI Fluoromount-G (Southern Biotech).

Images for ChR2 experiments in figures 3-5 were captured on an Olympus FV-3000 IX83 laser scanning confocal microscope including *Z* stack microscopy, while images in figure 6 were captured on an Olympus FluoView FV-1000 IX81 (thus resolution differences in IHC may be noted). Image capture was performed by sequential excitation of each fluorophore. Brain slices expressing GFP and mCherry from optogenetic experiments were mounted after cutting and imaged. To assess viral infection rates of ChR2-mCherry in GAD67-GFP positive cells and GAD65/GAD67/TH expression patterns, 4-8 higher magnification images (200-600x) were captured from each mouse for semi-quantitative analysis and n=4-5 mice were used for each experiment (see figure captions). Analysis was performed on Olympus cellSens software.

GAD67-GFP, ChR2-mCherry, GAD65-mCherry, TH-secondary antibody cellular staining was considered positive at greater than two standard deviations above background. Regions of interest were created based on significant fluorescence with size exclusion criteria to capture only intact full cells, which were then counted and confirmed by two separate individuals and averaged between all images from each mouse for n=1 per mouse.

### Acute and chronic cocaine treatment

Male and female GAD67 GFP^+^ mice (P20-35) or post-surgery VGAT::IRES-Cre/GAD67 GFP^+^ (P60-90) were intraperitoneally injected with cocaine (NIDA drug supply program) at 15 mg/kg either acutely (1 injection) or chronically (1 injection per day for 7 to 10 consecutive days) with the last dose 24 hours before sacrifice and experimentation.

### Brain slice preparation

Mice were anesthetized with isoflurane (1–2%) and decapitated with a rodent guillotine. Brains were rapidly removed and sectioned transversely on a vibratome at 230 μm (young adult) or 300 μm (juvenile). Juvenile brains were sliced using an ice-cold, sucrose-based cutting solution composed of 220 mM sucrose, 0.2 mM CaCl_2_, 3 mM KCl, 1.25 mM NaH_2_PO_4_, 25 mM NaHCO_3_, 12 mM MgSO_4_, 10 mM glucose, and 400 μM ascorbic acid. After sectioning, the slices were placed in oxygenated ACSF composed of 119 mM NaCl, 26 mM NaHCO_3_, 2.5 mM KCl, 1 mM NaH_2_PO_4_, 2.5 mM CaCl_2_, 1.3 mM MgSO_4_, and 11 mM glucose in an incubator at 35°C for an hour and then transferred to room temperature until recording. To better preserve adult VTA slices for electrophysiology recordings, adult brains were sliced using an ice-cold NMDG-based cutting solution composed of 92 mM NMDG, 2.5 mM KCl, 1.25 mM NaH_2_PO_4_, 30 mM NaHCO_3_, 20 mM HEPES, 25 mM glucose, 2 mM thiourea, 5 mM Na-ascorbate, 3 mM Na-pyruvate, 0.5 mM CaCl_2_·2H_2_O, and 10 mM MgSO_4_·7H_2_O, titrated to 7.5 pH with HCl. After sectioning, adult slices were kept in a warm bath of the NMDG-based cutting solution and were spiked with increasing amounts of NaCl over 20 min before being placed in an oxygenated HEPES holding solution composed of 92 mM NaCl, 2.5 mM KCl, 1.25 mM NaH_2_PO_4_, 30 mM NaHCO_3_, 20 mM HEPES, 25 mM glucose, 2 mM thiourea, 5 mM Na-ascorbate, 3 mM Na-pyruvate, 2 mM CaCl_2_·2H_2_O, and 10 mM MgSO_4_·7H_2_O titrated pH to 7.5 with concentrated 10 N NaOH (Ting et al., 2018).

### Electrophysiological recordings

Recordings began at least 1 h after cutting. Slices were placed in the recording chamber and bathed with oxygenated (95% O_2_, and 5% CO_2_) high divalent ACSF (4 mM CaCl_2_ and 4 mM MgSO_4_) at 30–32 degrees Celsius. The VTA was visualized using an Olympus BX51W1 microscope. GAD67/GFP^+^ cells were identified by fluorescence and located in the following coordinates from adult mouse bregma: anteroposterior −2.9 to −3.1, mediolateral 0.1 to 0.4, dorsoventral −3.9 to −4.1. GFP+ cells were patched with a borosilicate glass pipette (3–6 MΩ) filled with internal solution composed of 117 mM KCl, 2.8 mM NaCl, 20 mM HEPES, 5 mM MgCl_2_, 2 mM ATP-Na, 0.3 mM GTP-Na, and 1 mM QX-314 at pH 7.28 with an osmolarity at 275–285 mOsm. Recordings were made in voltage-clamp with GAD67^+^ or GAD65^+^/GAD67^-^ cells held at −65 mV. Prior data indicate GAD67 as an excellent marker for uniquely examining GABA cells as we and others noted previously (Chieng et al., 2011; Merrill et al., 2015).

Excitatory glutamate currents were blocked using 10µM CNQX (Alomone Labs) in non-optogenetic experiments. Plasticity was induced using an electrical 5 Hz stimulation in current clamp mode and delivered using a concentric bipolar electrode (Microprobes for Life Science) 200–300 µm from the patched cell. Currents were amplified using Multiclamp 700B and digitized with an Axon 1440A digitizer (Molecular Devices). Signals were filtered at 4kHz and recorded using Clampex 10.7 (Molecular Devices). Channelrhodopsin-induced synaptic currents were evoked by blue light (∼130 lux) generated by a TLED^+^ (Sutter Instruments) with individual light pulses lasting 0.5ms directed through the microscope lens. Note that in optogenetic experiments, evoked IPSCs were fully eliminated by GABA_A_ receptor antagonist, picrotoxin (100 µM). Cell input and series resistance were monitored continuously throughout each whole-cell experiment by 5 mV 100 ms hyperpolarization pulse; data were discarded if resistance changed by more than 15%.

### Drugs

CNQX (Alomone Labs), picrotoxin (Tocris) and cocaine (NIDA) were dissolved in ddH2O to make stock solutions. LY225910 (Tocris), N^6^CPA (Tocris) and SNAP (Tocris) were dissolved in DMSO to make stock solutions. CCK-8S (Tocris) was dissolved in PBS to make stock solutions. All stock solutions were frozen at –20/-80° C until diluted into ACSF to give the reported concentrations.

### Statistical Analysis

Electrophysiology data was analyzed and graphed using Clampfit software (Molecular Devices), Microsoft Excel, and Origin 10.8 (Origin Lab Corporation, Northampton, MA, USA). Statistical tests were computed using Microsoft Excel or SPSS. One-way ANOVA analysis was employed to determine if plasticity was statistically significant by comparing post-conditioning to baseline within every individual experiment. If an experiment was confirmed by ANOVA to display a significant (p < 0.05) increase in averaged post-conditioning responses compared to the baseline, it was grouped with iLTP; if the average post-conditioning responses showed a significant (p < 0.05) decrease compared to the baseline, the experiment was grouped with iLTD. Next, the average of the last 5 minutes of the baseline was compared in iLTD to 10–20 minutes post-conditioning, and for iLTP, to 20–30 minutes post-conditioning, as iLTP had a delayed onset to full plasticity induction (Nufer *et al*., 2023). Grouped data of either iLTP or iLTD was analyzed by ANOVA during these time points to confirm significance in post-conditioning responses to baseline. One-way ANOVA was also used to assess whether drug-induced changes in both individual and grouped experiments were significant by comparing the post-drug response to the last 5 minutes of the pre-drug response. Normality was assessed and if normally distributed, experiments were analyzed using student T-test (two-tail, unequal variance) to compare between two different experimental groups post-conditioning (i.e., control iLTP/iLTD to experimental iLTP/iLTD). For this analysis, the same 10-minute window at post-conditioning described above was picked from each group. To control multiple comparisons, a simple Bonferroni correction was implemented for electrophysiology data. A total of 27 comparisons for whole-cell experiments were performed, and a critical level for significance of 0.05/28 (0.0018) was used.

## Results

### iLTP/iLTD are CCK-independent

We previously identified that 5Hz low-frequency stimulation evoked either iLTP or iLTD of inhibitory inputs in different VTA GABA neurons. The iLTD was GABA_B_ receptor dependent, while the iLTP was partially NMDA receptor dependent (Nufer *et al*., 2023). We therefore further explored the mechanisms involved in iLTP. First, we examined CCK receptor involvement by applying CCK_2_R antagonist LY225910 (1 µM) while evoking the previously observed iLTP or iLTD. The presence of LY225910 did not alter either the iLTP or iLTD (Figure 1A). Interestingly, bath application of CCK-8S (0.1µM), the primary ligand of brain-expressed CCK_2_R, induced potentiation in ∼70% of VTA GABA neurons we examined (Figure 1B). To determine whether the CCK-induced potentiation occludes the 5Hz-evoked iLTP, we applied the 5Hz stimulus at the end of an established CCK-induced potentiation (Figure 1C-E).

**Figure 1.**
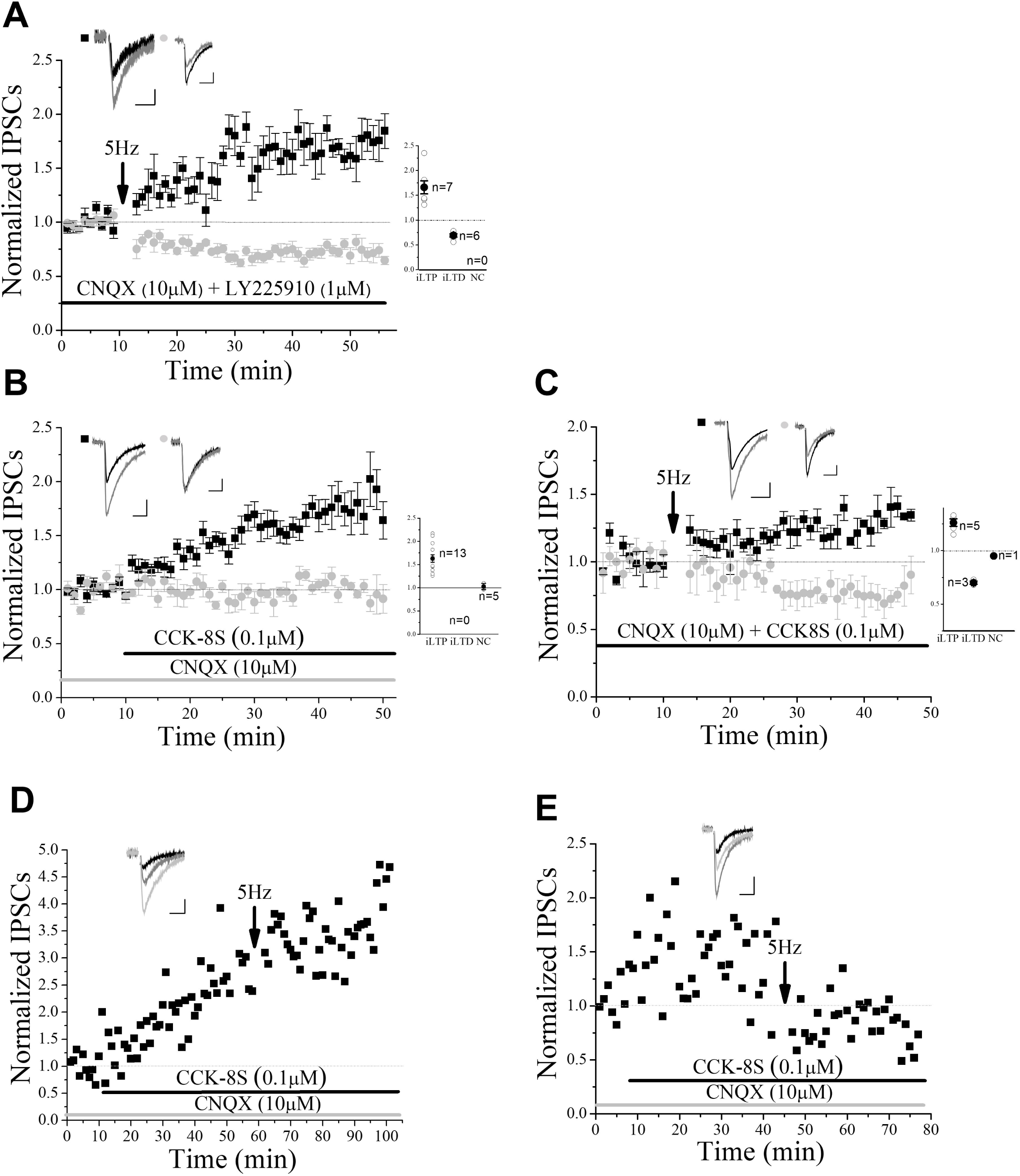
CCK potentiates inhibitory currents but does not mediate iLTP. A) Bath application of CCK2R antagonist LY225910 (1 µM) had no impact on iLTP or iLTD induced by a 5-Hz (arrow) conditioning stimulus (iLTP: p<0.0001, compared to baseline, ANOVA, n=7; iLTD: p<0.0001 compared to baseline, ANOVA, n=6) as measured by normalized GABAergic inhibitory postsynaptic currents (IPSCs). B) Bath application of CCK receptor agonist, CCK-8S (0.1µM), potentiated 72% of the ventral tegmental area (VTA) GABA neurons examined (Potentiation: p<0.0001, compared to baseline, ANOVA, n=13; no change: p>0.05, compared to baseline, ANOVA, n=5). C) Occlusion experiments using CCK8S pre-exposure did not alter iLTP or iLTD (iLTP: p<0.0001, compared to baseline, ANOVA, n=5; iLTD: p<0.0001 compared to baseline, ANOVA, n=4). D) A representative experiment from (C) including baseline prior to CCK8S exposure, illustrates iLTP following 5Hz stimulation occurs following CCK8S-induced potentiation of IPSCs. E) A similar representative experiment also demonstrates following CCK8S-induced potentiation that iLTD continues to occur as well. IPSC trace insets from A-C: baseline (black), 5Hz-or CCK-induced changes (grey). Inset IPSC traces from D-E: baseline (black), CCK-8S application (grey), 5Hz-induced iLTP or iLTD (light grey). Scale bars for traces here and throughout are 100 pA and 10 ms.

Neither iLTP nor iLTD were occluded, collectively suggesting CCK receptors are present at most GABAergic inputs to VTA GABA cells, but are not involved in either plasticity.

### Nitric oxide is not required for iLTP

When we investigated the possible involvement of the nitric oxide (NO) pathway in the observed iLTP previously, we noted neither iLTP nor iLTD was altered by blocking NO production, however the NO donor SNAP potentiated just over half of the GABA neurons we examined (Nufer *et al*., 2023). To follow up on this finding we conducted an occlusion experiment by applying the 5Hz stimulus at the end of a SNAP-induced potentiation (Figure 2). In this case, SNAP occluded iLTP suggesting a possible common signaling cascade for iLTP induction and NO.

**Figure 2.**
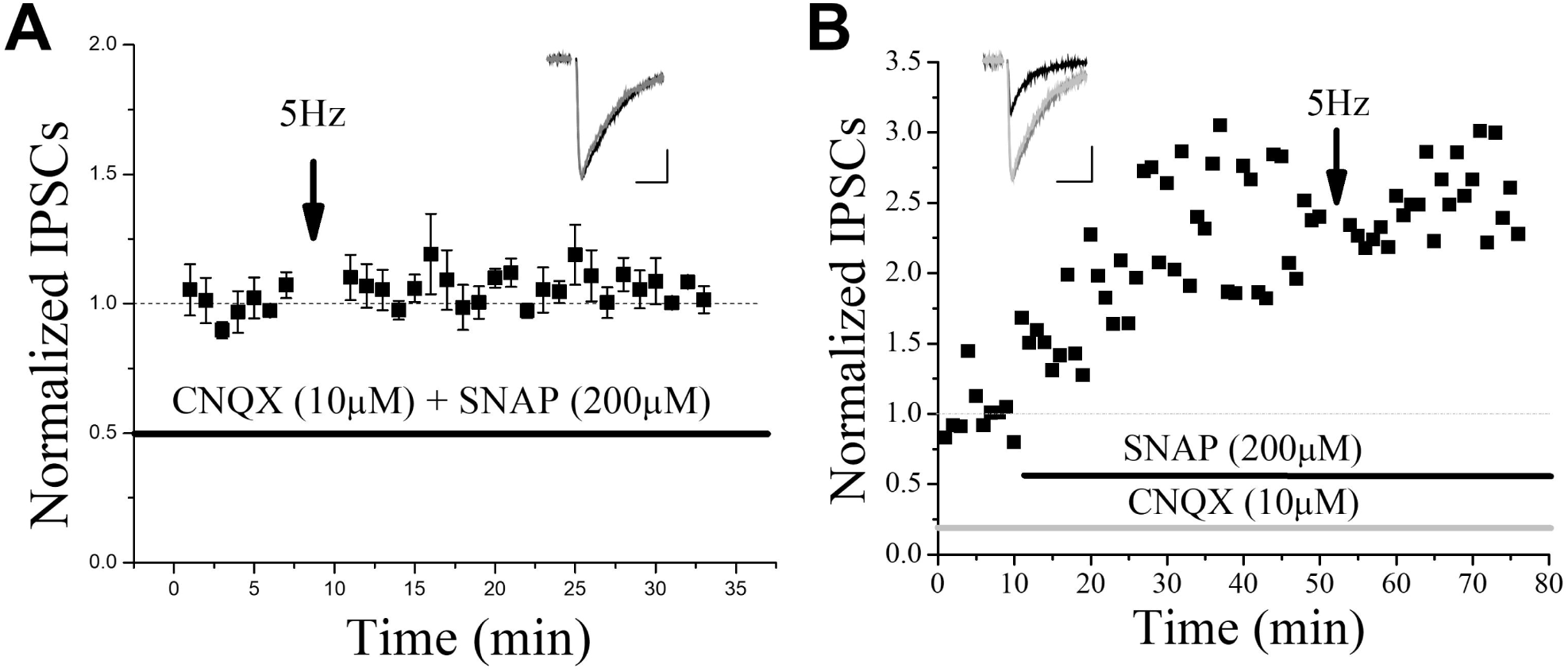
The nitric oxide pathway likely signals through the same pathway inducing iLTP. A) Nitric oxide donor SNAP (200 µM) occluded iLTP induction by the 5Hz stimulus (occlusion: p=0.09, compared to baseline, ANOVA, n=4). B) A representative experiment including baseline prior to SNAP application illustrates SNAP-induced potentiation, and occlusion of 5Hz-induced iLTP. Inset IPSC traces in A: baseline (black), 5Hz-induced changes (grey); B: baseline (black), SNAP application (grey), 5Hz-induced changes (light grey).

**Figure 3.**
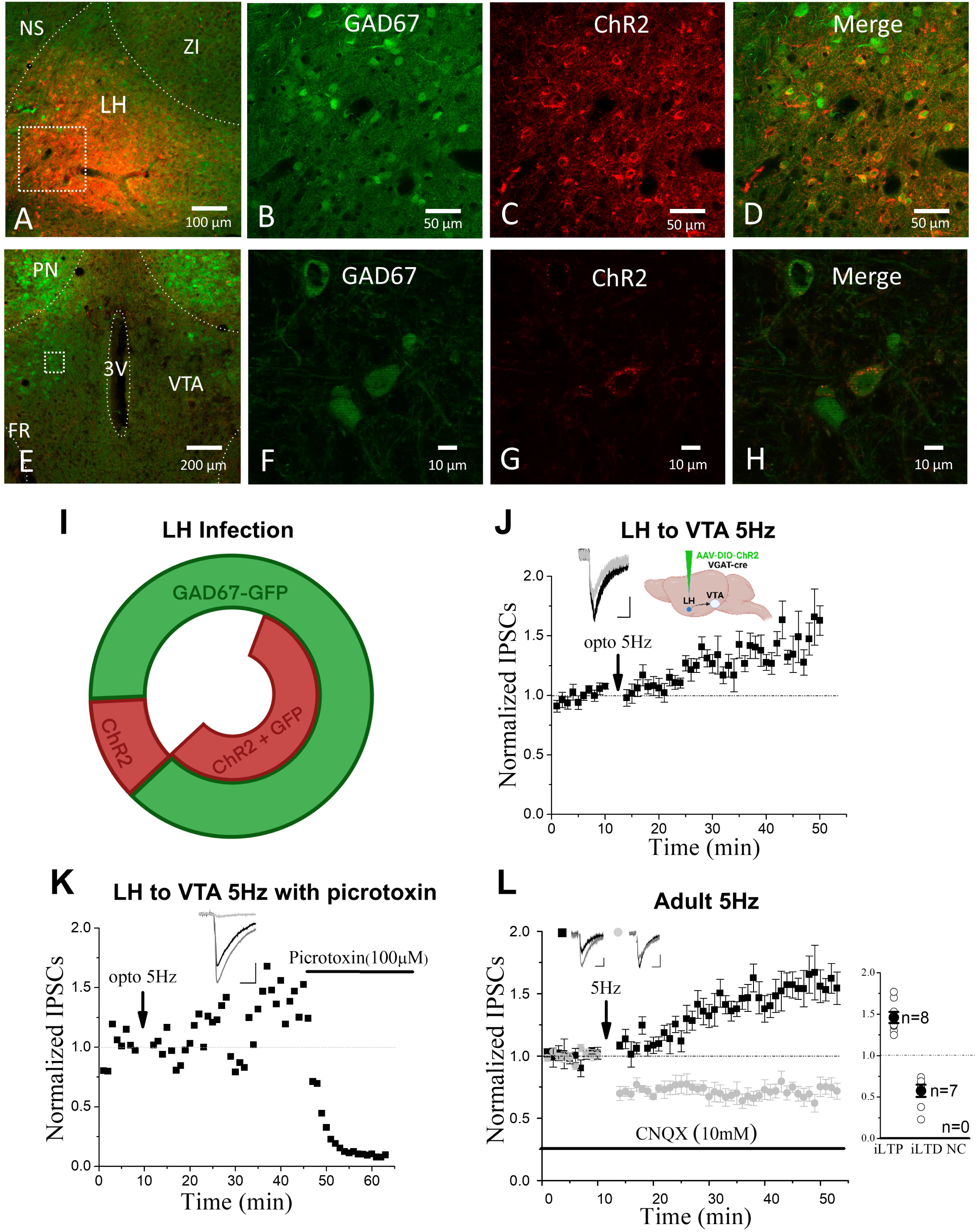
Optogenetic stimulation of lateral hypothalamus (LH) inhibitory inputs to VTA GABA cells selectively evoked iLTP. A) Immunohistochemistry (IHC) of a 100x overlay demonstrates the LH surgical injection location of a Cre-inducible channelrhodopsin (ChR2) containing virus in VGAT::IRES-Cre/GAD67 GFP^+^ mice. The white square highlights the location of 400x images of GAD67-GFP (B) and ChR2-mCherry (C) along with the overlay of both (D), demonstrating ChR2 expression in GAD67-positive GABA cells in the LH. E) A 100x IHC image illustrates the VTA. The white square in E is where 600x images are taken demonstrating GAD67 GABA cells in the VTA (F), surrounded by ChR2-containing puncta originating from axonal inputs from infected ChR2-mCherry containing LH neurons (H, overlay). I) Semi-quantitative analysis of IHC demonstrates 58.7 ± 8.5% of GAD67-expressing neurons in the LH co-express GAD67-GFP and ChR2-mCherry after viral infection, while 14.2 ± 4% of ChR2 infected cells did not co-express GAD67 GFP (n=5 mice). The majority of the latter cells are likely GAD65-only expressing GABAergic cells as we employed Vgat-Cre mice to target all GABA cells but used a GAD67 marker in order to positively identify unique GABA cells in the VTA (see figure 6 where some GAD65 cells also express TH). J) The mCherry decorated GAD67^+^ cells in the VTA were recorded using whole-cell voltage clamp electrophysiologically and when LH inputs were optogenetically stimulated at 5Hz (0.5ms duration pulses) they selectively induced iLTP only (iLTP: p<0.0001 compared to baseline, ANOVA, n=13). Inset cartoon indicates injection location of ChR2-mCherry containing virus. K) A representative experiment from J demonstrates optically induced iLTP, which was eliminated by GABA_A_ receptor antagonist picrotoxin (100 µM), indicating all LH inputs recorded from in the VTA were indeed GABAergic, and thus confirm the vast majority of LH infected mCherry^+^/GAD67^-^ cells were GABAergic (GAD65-only). Note: CNQX was not applied during optogenetically evoked IPSC experiments. L) Adult mice also exhibit both iLTP and iLTD that we previously observed in juvenile/adolescent animals at the same ∼50/50 ratio, which were not significantly different from these younger mice (iLTP: p<0.0001, compared to baseline, ANOVA, n=8, and P=0.2380 compared to (Figure 1; (Nufer *et al*., 2023)), Student’s T-Test; iLTD: p<0.0001 compared to baseline, ANOVA, n=7, and P=0.2457 compared to (Nufer *et al*., 2023), Figure 1, Student’s T-Test). Note the delay plasticity onset, particularly of iLTP (J, L) are consistent with prior published data in adolescent mice (Nufer *et al*., 2023). Inset IPSC traces in K: baseline (black), 5Hz-induced changes (grey); L: baseline (black), optogenetic 5Hz-induced iLTP (grey), picrotoxin application (light grey). Abbreviations: NS, nigrostriatal bundle; LH, lateral hypothalamus; ZI, zona incerta; PN, Posterior Hypothalamic Nucleus; 3V, third ventricle; FR, fasciculus retroflexus.

### Plasticity at LH inhibitory inputs to VTA GABA cells

Next, to examine the hypothesis that regional selective GABA input specificity is responsible for this rare bipotential plasticity phenomenon, we employed optogenetics using stereotaxic selective surgical injection of a Cre-inducible AAV-ChR2 virus in a Vgat-Cre mouse line. The first input source we examine was LH as it sends GABA inputs to VTA GABA neurons (Faget *et al*., 2016). We injected the LH of VGAT::IRES-Cre/GAD67 GFP^+^ mice with AAV2/1-EF1a-DIO-HhChR2(H134R)-mCherry. Post-surgery IHC images confirmed that ChR2-mCherry^+^ cells co-expressed at a high percentage of GAD67^+^-GFP GABA cells in the LH (Figure 3A-D, I). In addition, we noted that mCherry^+^ axon terminals from the LH localized near GAD67/GFP^+^ neurons in the VTA (Figure 3E-H), which were visually selected for whole cell electrophysiology experiments. In response to an optical 5Hz stimulation following the baseline, inhibitory inputs from LH GABA to VTA GABA neurons induced exclusively iLTP (Figure 3J). This optically induced response was GABA_A_-mediated, as bath application of picrotoxin eliminated the IPSCs (Figure 3K). To control for the age influence on plasticity induction, we performed experiments in adult mice, given that the subjects in the optogenetic experiments were typically young adults aged 60 to 90 days postnatal due to post-surgery time required for ChR2 expression. Indeed, 5Hz conditioning stimulus continued to induce either iLTP or iLTD in adult mice (Figure 3L) to the same degree as in juvenile/adolescent animals (Nufer *et al*., 2023).

### Plasticity at RMTg inhibitory inputs to VTA GABA cells

The RMTg also sends GABA inputs to VTA GABA neurons in addition to VTA DA cells where the latter leads to DA cell disinhibition (Jhou *et al*., 2009; Kaufling et al., 2010).

VGAT::IRES-Cre/GAD67-GFP mice received injections of AAV2/1-EF1a-DIO-HhChR2(H134R)-mCherry into the RMTg. IHC images following surgeries indeed indicate high expression of ChR2-mCherry^+^ positive in GAD67-GFP^+^ cells within the RMTg (Figure 4A-E).

**Figure 4.**
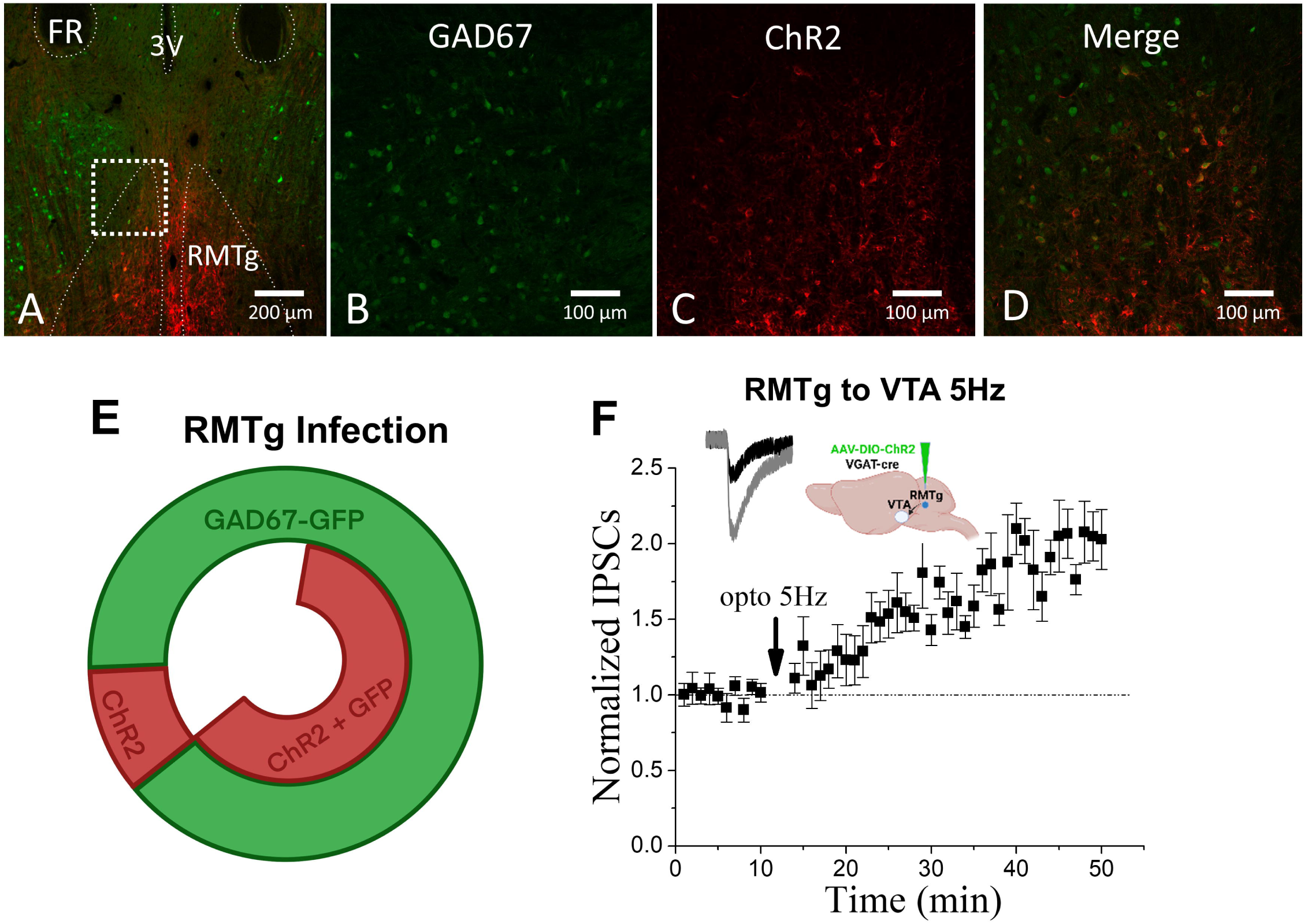
Optogenetic stimulation of rostromedial tegmental nucleus (RMTg) inhibitory inputs to VTA GABA cells selectively evoked iLTP. Experimental methods and mouse models were the same as in figure 3 except the surgical injection site was the RMTg. A) IHC of a 100x overlay demonstrates the RMTg surgical injection location of a Cre-inducible ChR2-containing virus. The white square highlights the location of 200x images of GAD67-GFP (B) and ChR2-mCherry (C) along with the overlay (D), demonstrating ChR2 expression in RMTg GAD67-positive GABA cells. E) Semi-quantitative analysis of IHC demonstrates 66.8 ± 1.5% of GAD67-expressing neurons in the RMTg co-express GAD67-GFP and ChR2-mCherry after viral infection, while 10.2 ± 2.6% of ChR2-mCherry infected cells did not co-express GAD67 GFP (n=4 mice). F) The mCherry decorated GAD67^+^ cells in the VTA were recorded using whole-cell voltage clamp electrophysiologically and when RMTg inputs were optogenetically stimulated at 5Hz they selectively induced iLTP (iLTP: p<0.0001 compared to baseline, ANOVA, n=9). Inset cartoon indicates injection location of ChR2-mCherry containing virus. Inset IPSC traces in F: baseline (black), optogentic 5Hz stimulus induced iLTP (grey).

**Figure 5.**
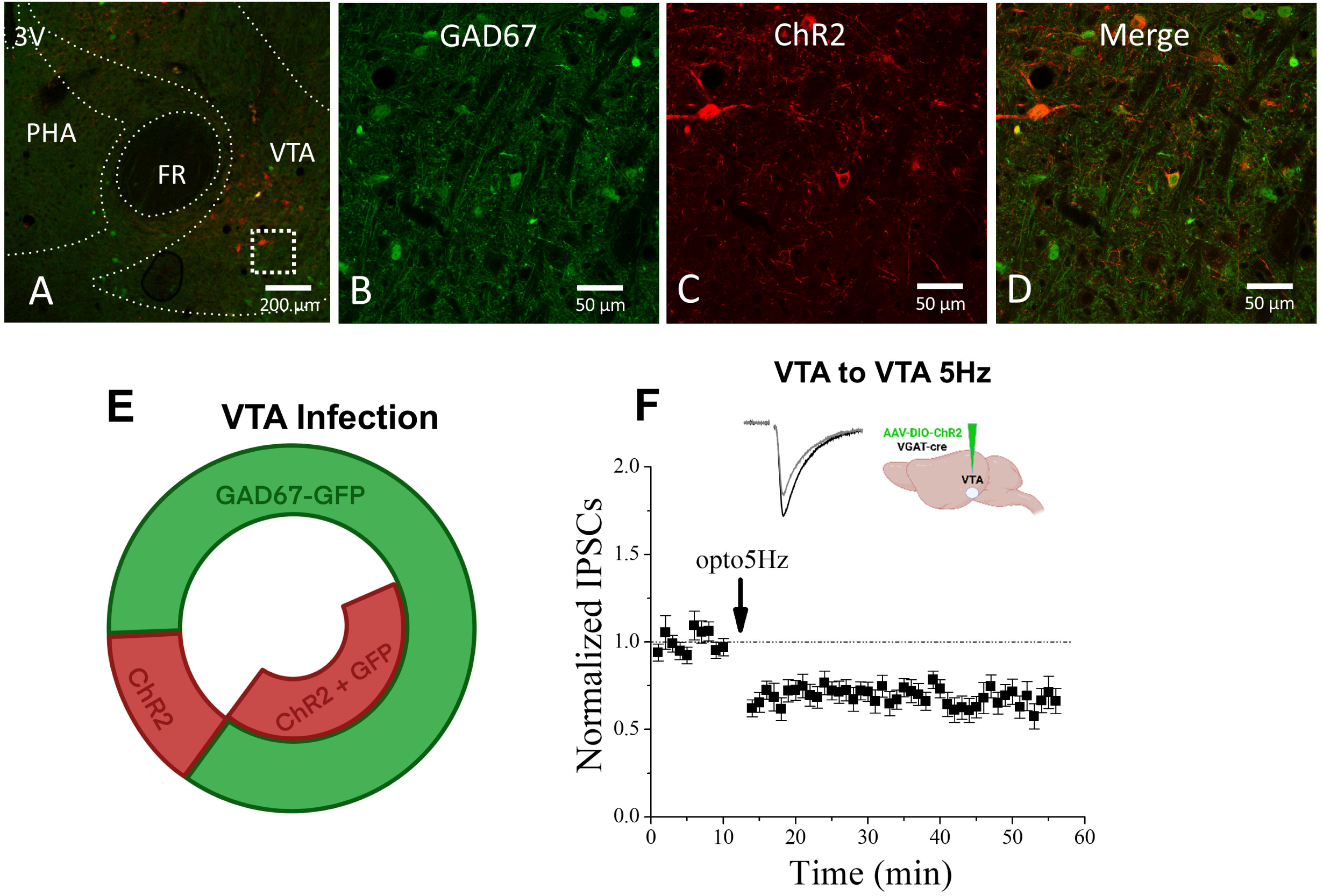
Optogenetic stimulation of local VTA inhibitory inputs to VTA GABA cells selectively evoked iLTD. A) IHC of a 100x overlay demonstrates the VTA surgical injection location of a ChR2-mCherry containing virus (500 nL). The white square highlights the location of 400x images of GAD67-GFP (B) and ChR2-mCherry (C) along with the overlay (D), demonstrating ChR2 expression in some, but not all GAD67-positive VTA GABA cells. E) Semi-quantitative analysis of IHC demonstrates 41.8 ± 3.4% of GAD67-expressing neurons in the VTA co-express GAD67 and ChR2-mCherry after viral infection, while 14.4 ± 3.1% of ChR2 infected cells did not co-express GAD67 GFP (n=5 mice). F) Non-mCherry labeled GAD67^+^ VTA cells were recorded using whole-cell voltage clamp electrophysiologically where 5Hz optogenetic stimulation of local VTA GABA inputs selectively induced iLTD (iLTD: p<0.0001 compared to baseline, ANOVA, n=11). Inset cartoon indicates injection location of ChR2-mCherry containing virus. Inset IPSC traces in F: Baseline (black), optogentic 5Hz stimulus induced iLTD (grey). Abbreviation: PHA, Posterior Hypothalamic Area.

In response to an optical 5Hz stimulation, inhibitory inputs from RMTg to VTA GABA neurons decorated with mCherry also induced exclusively robust iLTP (Figure 4F). This iLTP is significantly larger (21.3±1.8%; P<0.0001; Student T test) than iLTP observed at the LH to VTA GABA synapses and had a faster onset.

### Plasticity at local VTA inhibitory inputs to VTA GABA cells

Within the VTA, GABA neurons not only send direct contact to the neighboring dopamine neurons but also form local connections with each other (Dobi et al., 2010; Omelchenko and Sesack, 2009). AAV2/1-EF1a-DIO-HhChR2(H134R)-mCherry was injected into the VTA of VGAT::IRES-Cre/GAD67-GFP^+^ mice. Post-surgery IHC images confirmed a high level of successful transfection (Figure 5A-E). We avoided recording from VTA cells that were directly infected by ChR2-mCherry (i.e., strong Na^+^ spikes would be present if patched) and only selected non-infected GFP^+^ GABA neurons that received innervation from infected cells (i.e., mCherry puncta near cell soma). In response to an optical 5Hz stimulation, the intrinsic VTA inhibitory inputs induced exclusively iLTD in all VTA GABA neurons examined (Figure 5F).

### Plasticity displayed by VTA GAD65-only neurons

An additional potential explanation for the bipotential plasticity phenomenon observed at GABA inputs to VTA GABA cells is the existence of subtypes of VTA GABA neurons giving rise to disparate forms of plasticity. Our approach to investigating this hypothesis was by examining the distribution of GABA synthesis enzymes GAD67 and GAD65 in VTA neurons. We examined GAD67/GFP^+^-GAD65/mCherry^+^ crossed mice to visualize simultaneously the VTA GAD67 and GAD65 expression. We performed immunohistochemistry on fixed 25-30 µM transverse cryostat cut slices in the VTA (Figure 6 A, B) using anti-tyrosine hydroxylase (TH) primary antibody and blue-fluorescing secondary antibody. TH (Figure 6 C, D), GAD67 (Figure 6 E, F) and GAD65 (G, M) are independently illustrated along with the overlay of all three (Figure 6 I, J). We noted three major types of GAD expression including GAD65-only, GAD67-only, and the majority cell type GAD65/GAD67 co-expressing cells (Figure 6 K). We also noted some TH-positive dopamine cells also co-express GAD65 (Figure 6 L; see figure 6 I, J). Near the VTA midline we saw a consistent population of GAD65-only expressing cells by themselves (Figure 6 M, N). Prior reports have noted unique expression patterns of DA cell subtypes in different quadrants of the VTA, usually distributed medially to laterally (Brischoux et al., 2009; Stamatakis et al., 2013). Similarly, we note groupings of putative subtypes of cells including GABA cells. For example, figure J illustrates a group of DA neurons that nearly all express GAD65, even though this is a minority cell subpopulation, along with GAD67 cell groupings above and below the TH-positive DA cells. Note the lack of GAD67 and TH co-expression in these images again indicating GAD67 as an excellent marker of VTA GABA cells selectively. In addition, GAD65-only cells also group together in medial VTA. While we did not perform a full quantification of cellular subtype by location, which was not the intent of this study, we reported potential subpopulation groupings and illustrate the presence of GAD65-only expressing cells (see Figure 6 M, N). This allowed us to evaluate our hypothesis of differential plasticity on a unique GABA cell population, as all our studies to this point were on GAD67-GFP^+^ cells. We utilized GAD67-GFP^+^/GAD65-mCherry^+^ mice for whole cell electrophysiology experiments and targeted GAD65-only cells (GFP^-^/mCherry^+^) in the same location where GAD65 did not overlap with TH. Data from these cells exhibited the same bipotential plasticity phenomenon (Figure 6 O), which we noted in GAD67-GFP^+^ cells here (see Figure 3L) and in our previous study (Nufer *et al*., 2023), in a similar 50-50 ratio. Interestingly, unlike what we saw in GAD67-expressing cells (Nufer *et al*., 2023), which all have excitatory inputs like DA cells, the application of CNQX caused a 48±1.1% reduction of EPSCs in only half of the GAD65-only cells examined (Figure 6P), indicating that a portion of GAD65-only cells do not receive glutamatergic inputs, and likely make up a novel GABA cell subtype in the VTA.

### Cocaine eliminates iLTD

As cocaine modification of DA cell synaptic plasticity is a hallmark of dependence induction, its impact on GABA cell synaptic plasticity was essential to examine. We therefore conducted whole-cell electrophysiology experiments in animals receiving either acute or chronic IP injection of cocaine. Both chronic and acute cocaine treatment of GAD67 GFP^+^ mice eliminated iLTD but not iLTP at GABA inputs to VTA GABA cells (Figure 7 A, B). Control animals receiving saline IP injections for 7 to 10 consecutive days continued to display both iLTP or iLTD as usual in response to 5Hz stimulation (Figure 7C). Since optogenetic stimulation at VTA to VTA GABA synapses evoked exclusively iLTD (Figure 5F), we examine whether this opto-induced iLTD was also sensitive to cocaine exposure. Chronic cocaine was administered to VGAT::IRES-Cre/GAD67 GFP mice that received injection of AAV2/1-EF1a-DIO-HhChR2(H134R)-mCherry in the VTA. As expected, iLTD in VTA GABA neurons was prevented by chronic cocaine injection (Figure 7D). It was previously demonstrated that chronic cocaine treatment can modify DA receptor D_1_ (D_1_R) regulation of GABA_B_ IPSCs through augmented adenosine tone in VTA DA neurons (Bonci and Williams, 1996). Also, D1-containing GABA afferents sent from NAc to VTA are modulated by adenosine A_1_ receptors (A_1_R) (Edwards *et al*., 2017). Combining literature evidence with the observation that iLTD is GABA_B_R dependent (Nufer *et al*., 2023) as well as the confirmed presynaptic presence of A_1_R at all VTA GABA afferents tested (Figure 7E), we hypothesized that the loss of iLTD after cocaine treatment was caused by increased adenosine tone and could be recovered by blocking A_1_R activation. Using chronic cocaine treated GAD67 GFP^+^ animals with bath applied DPCPX (1 µM), an A_1_R antagonist, iLTD in VTA GABA cells is recovered (Figure 7F).

**Figure 6.**
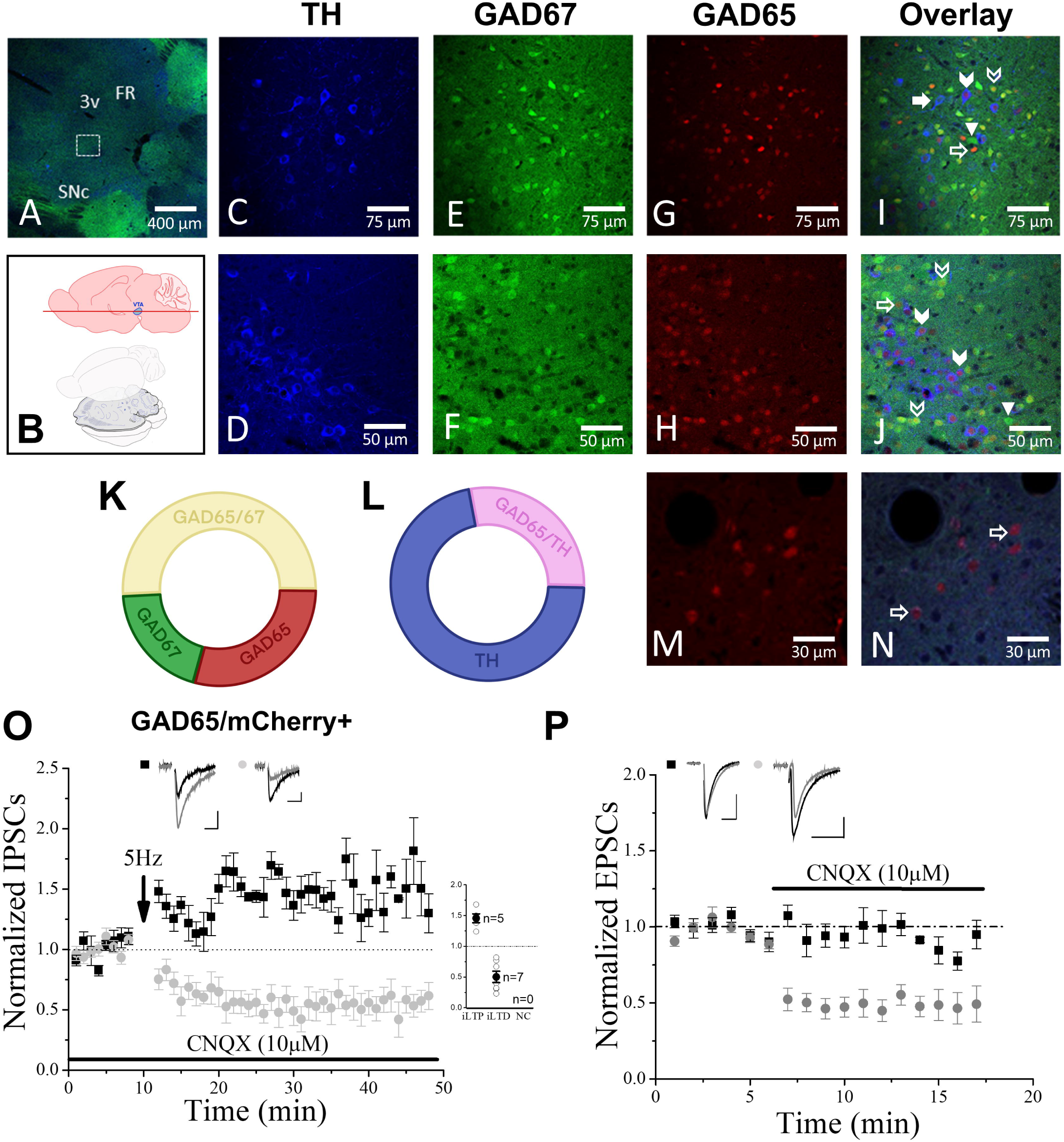
Heterogeneity of GAD65, GAD67, and tyrosine hydroxylase (TH) expression among VTA GABA and dopamine neurons. A) Low magnification (50x) IHC image highlights one of the general VTA areas where images were selected for analysis. B) Schematic diagram demonstrates the level of transverse slices made using frozen cryostat sections to illustrate VTA cell types. TH (C, D), GAD67-GFP (E, F) and GAD65-mCherry (G, H) IHC 400x images taken from two different mice are used to highlight unique expression of these three targets in different VTA cells demonstrated by the overlay of all three IHC images (I, J). Cells highlighted include GAD67-only (arrowhead), GAD65-only (open arrow), GAD67/GAD65 co-expressing (open chevron), TH-only (filled arrow), and TH/GAD65 co-expressing (filled chevron) cells. The presence of TH in IHC images highlights that we are indeed examining the VTA as the location of analyzed images. K) The pie chart illustrates percentages of VTA GABAergic cell categorizations identified as GAD67/GAD65 co-expressing cells (50.9%) that make up most GABAergic cells, GAD65-only cells (29.1%), and GAD67-only cells (20%). L) The pie chart illustrates TH-only positive cells (71.6%) were the most expressed dopaminergic cell, with a smaller percentage of TH-positive/GAD65 co-expressing cells (28.4%). As highlighted by others, GABA or DA cells of similar type tend to cluster (i.e. see a high percentage of the rarer TH-positive/GAD65 co-expressing cells in image J). Importantly, only rare (<<1%) co-expression of GAD67 (GFP) and TH (blue stain) was identified in the VTA (only a few cells). Note also that these expression percentages in K and L are not likely representative of the entire VTA, as we focused image capture and analysis on regions of highest GAD67/GAD65 expression. M) GAD65-only cells are located in highest concentration at the midline of the VTA in our analysis (400x) and are clearly TH– and GAD67-negative in the overlay (N). O) Averaged IPSC data from these GAD65-only cells using whole-cell voltage clamp recordings illustrate that 5Hz stimulation induces bipotential plasticity of iLTP and iLTD (iLTP: p<0.0001, compared to baseline, ANOVA, n=5; iLTD: p<0.0001 compared to baseline, ANOVA, n=8), which we observed in GAD67-positive cells here (see figure 3 L) and previously (Nufer *et al*., 2023). P) Bath application of CNQX caused a 48±1.1% reduction of IPSCs in half of the GAD65-only cells examined (depression: p<0.0001, compared to baseline, ANOVA, n=6; no change: p=0.1282 compared to baseline, ANOVA, n=7). Inset IPSC traces in baseline (black), 5Hz-induced changes or CNQX application (grey); P: baseline (black), CNQX application (grey). Abbreviations: SNc, substantia nigra; 3v, third ventricle; FR, fasciculus retroflexus.

**Figure 7.**
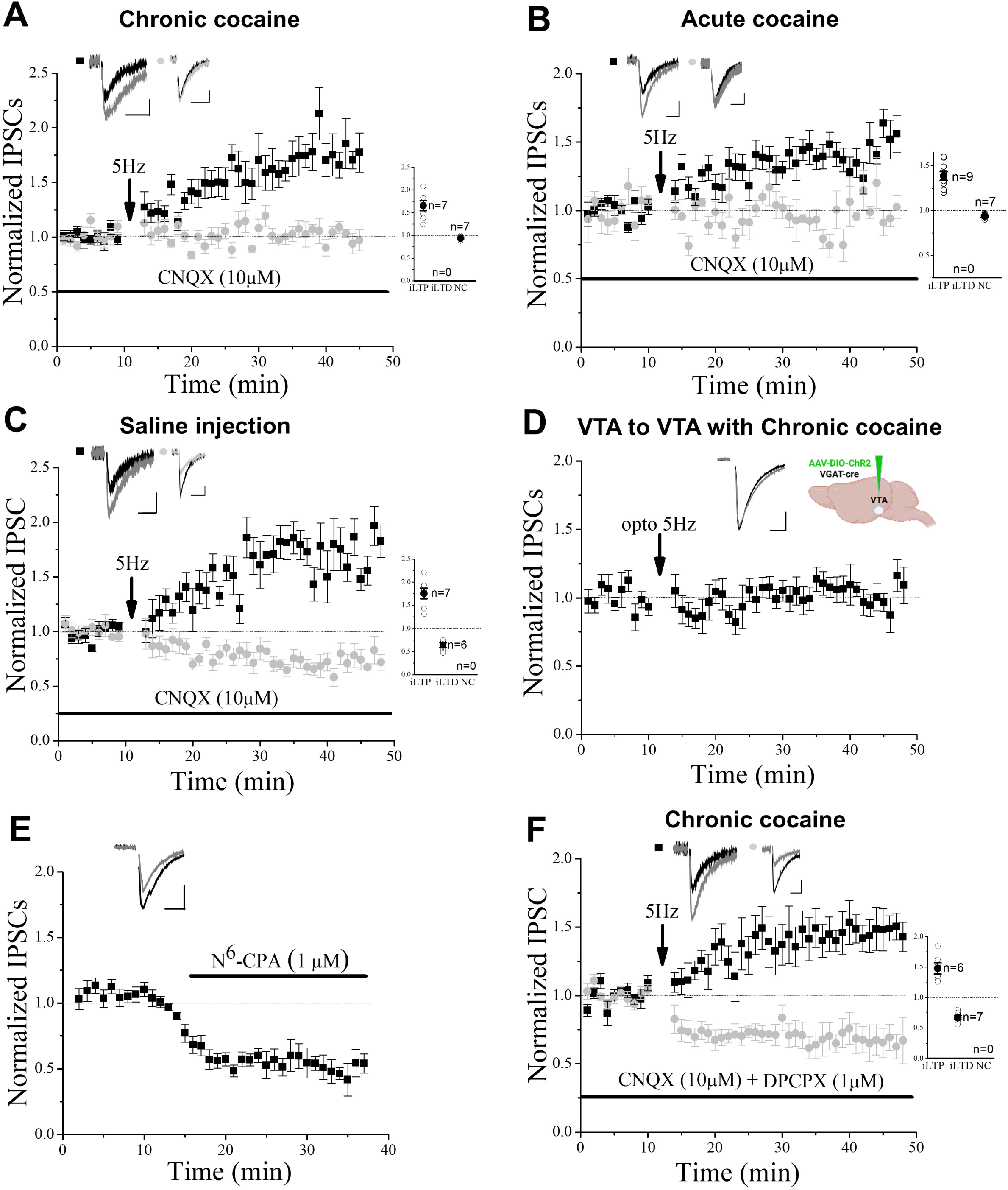
Cocaine treatment eliminated and electrically evoked iLTD, and optically evoked iLTD from local VTA GABA inputs to VTA GABA cells. A) Chronic cocaine treatment (7-10 days, daily intraperitoneal injections) prevented the 5Hz-induced iLTD (no plasticity group: p=0.1937, compared to baseline, ANOVA, n=7, and p<0.0001 compared to control iLTD from figure 5M or our prior study (Nufer *et al*., 2023), Student’s T-Test); iLTP: p<0.0001, compared to baseline, ANOVA, n=7). B) Acute cocaine treatment also prevented the 5Hz-induced iLTD (no plasticity group: p=0.8362, compared to baseline, ANOVA, n=7, and P<0.0001 compared to control iLTD from figure 5M or our prior study (Nufer *et al*., 2023), Student’s T-Test; iLTP: p<0.0001, compared to baseline, ANOVA, n=9). C) Saline-injected (7-10 days, daily intraperitoneal injections) mice displayed both 5Hz-induced iLTP or iLTD (iLTP: p<0.0001, compared to baseline, ANOVA, n=7; iLTD: P<0.0001, compared to baseline, ANOVA, n=6). D) Chronic cocaine treatment prevented iLTD evoked by optogenetic stimulation at the local VTA GABA inputs to VTA GABA cells (no plasticity group: P=0.4046, compared to baseline, ANOVA, n=6, and P<0.00001, compared to control optogenetically induced 5Hz-evoked iLTD in figure 5F, Student’s T-Test). Inset cartoon indicates injection location of ChR2-mCherry containing virus. E) Activation of the adenosine A1 receptor (A_1_R) by adenosine agonist N^6^-CPA application caused a 47±3.4% reduction of IPSCs in all VTA GABA neurons tested (depression: p<0.0001, compared to baseline, ANOVA, n=12). F) Bath application of A_1_R antagonist DPCPX (1 µM) restored iLTD in chronic cocaine treated animals (iLTP: p<0.0001, compared to baseline, ANOVA, n=6; iLTD: P<0.0001, compared to baseline, ANOVA, n=7). Inset IPSC traces: baseline (black), 5Hz induced changes or N^6^CPA application (grey).

## Discussion

Growing evidence of VTA GABA cell involvement in reward processes (Bouarab et al., 2019; Creed *et al*., 2014) including self-stimulation (Lassen *et al*., 2007), conditioned place aversion (Tan *et al*., 2012), conditioned place preference (Bocklisch *et al*., 2013), and associate reward learning (Brown *et al*., 2012), necessitated further examination of these cells and their plasticity. Previously, we observed a unique bipotential plasticity (iLTP or iLTD) of inhibitory inputs to VTA GABA cells, where iLTD was GABA_B_R dependent and iLTP displayed partial NMDA receptor-dependence (Nufer *et al*., 2023). Here we determined using optogenetic activation that induction of iLTP or iLTD is synapse-input specific, and that cocaine exposure eliminates iLTD, while sparing iLTP. This latter finding combined with our prior demonstration of alcohol elimination of only iLTD (Nufer *et al*., 2023) suggests that iLTD of GABA inputs to VTA GABA cells is potentially a crucial target of addictive substances maladaptive VTA synaptic modulation of the inhibitory circuit.

### CCK is not involved in iLTP

CCK receptors participate in hippocampal and cortical LTP (Chen et al., 2019; Su et al., 2023), where CCK restores morphine-attenuated LTP in rat dentate gyrus (Wen et al., 2014). In the VTA, Damonte et al. (2023) reported that somatodendritic release of CCK from VTA DA neurons induced presynaptic LTP_GABA_ (i.e., GABA input to DA cells) via CCK_2_ receptors (Martinez Damonte *et al*., 2023). In the current study while the predominant CNS CCK isoform (Crawley, 1985; Rehfeld, 2017), CCK-8S, induced a potentiation in over half of inhibitory VTA GABA neurons, indicating synaptic CCK receptor expression, whether these are the same cells exhibiting iLTP we cannot say, but it is potentially the case. CCK-8, however, failed to occlude either iLTP or iLTD, suggesting that while the CCK receptor is present, it is not involved in either plasticity, unlike VTA DA cell iLTP. The presence of CCK receptors at differential inputs demonstrates additional complexity of the GABAergic circuitry in the reward system as it modifies some, but not all inputs to VTA DA and GABA cells.

### NO is not required for iLTP

A previous study identified that the GABAergic LTP (i.e. iLTP or LTP_GABA_) on VTA DA cells was NO-dependent (Nugent *et al*., 2007). In contrast, we previously noted that nNOS inhibitors did not alter iLTP/iLTD of VTA GABA cells, though we did note that SNAP (NO donor) potentiated ∼50% of VTA GABA neurons (Nufer *et al*., 2023), similar to GABA inputs to VTA DA neurons (Nugent *et al*., 2007). Therefore, we re-examined SNAP noting it did indeed occlude iLTP, suggesting that while iLTP is independent of nNOS, it likely requires a similar downstream signaling pathway. This potential signaling mechanism is currently being examined.

### Bipotential plasticity response is likely input-specific

The occurrence of LTP/iLTP or LTD/iLTD with the exact same stimulus parameters in a bipotential manner has rarely been reported in existing literature, and never in GABAergic cells that we are aware. In our previous study, we proposed the hypotheses that plasticity type could be input selective or based on GABA cell sub-populations (Nufer *et al*., 2023). To selectively activate regional inputs, we employed optogenetics, which illustrated local GABA inputs to VTA GABA cells exhibit iLTD, while GABA inputs projecting from RMTg and LH exhibited iLTP, suggesting plasticity type is indeed input specific. Recent similar finds by others on VTA DA cells illustrated their inhibitory inputs also exhibit a similar bipotential plasticity phenomenon that was input selective (St Laurent and Kauer, 2019). In addition, as activation of inhibitory inputs selectively from the NAc to VTA GABA cells only induce iLTP (Bocklisch *et al*., 2013), further supports this hypothesis. Lastly, our data highlight the possibility that ∼50% of VTA GABA cells receive mainly local GABA connections, while the other 50% are innervated by afferent GABA inputs from outside the VTA, or at least a preponderance of one or the other, based on the ∼50% split of iLTD or iLTP exhibition in VTA GABA cells (Nufer *et al*., 2023).

Lastly, a methodological caveat for our optogenetic studies here is that the RMTg and VTA are extremely close, and viral injection of one region could have infected the other. However, IHC images suggest good localization of ChR2-containing viral infection, and iLTD only occurred in VTA injections and iLTP in RMTg injections, suggesting good ChR2 targeting.

Our second hypothesis of correlating plasticity to GABA cell subtype based on GAD65/67 expression seems less plausible. The observation that GAD65-only cells also displayed bipotential plasticity response, akin to GAD67-expessing cells, suggests plasticity type is not distinguished by the distribution of GABA synthesis enzymes GAD67 and GAD65 in VTA GABA neurons. However, the complexity of the VTA’s heterogeneity, as revealed by recent molecular studies (Paul *et al*., 2019; Phillips *et al*., 2022), suggests GABA subtypes characterized using other markers would be valuable for future examination. Therefore, it would be imprudent to dismiss the cell subtype hypothesis outright without further investigation, which we are currently investigating further. An important note regarding GAD enzymes is that some DA neurons co-release other neurotransmitters including GABA, via co-expression of TH and GAD65, but never GAD67 (Barker et al., 2016; Gonzalez-Hernandez et al., 2001; Merrill *et al*., 2015; Morales and Margolis, 2017; Olson and Nestler, 2007; Phillips *et al*., 2022; Stamatakis *et al*., 2013). We note here that both GAD65^+^/TH^+^ co-expressing, and GAD65-only neurons are present in medial VTA, but the latter are situated more dorsally. During electrophysiology experiments, we identified and recorded medial-dorsal GAD65-only neurons (mCherry^+^/GFP^-^). As the mCherry^+^/GFP^-^ cells we examined still exhibit either iLTP or iLTD in a similar ratio as GAD67^+^-only (GFP) cells, and approximately half of the GAD65^+^-only cells we recorded receive minimal to no glutamate inputs, unlike DA cells, this suggests that these cells are likely a functionally novel GABA cell subtype. Indeed, the lack of synaptically-evoked glutamate currents onto some GAD65^+^-only cells, in contrast to GAD67^+^ cells (Nufer *et al*., 2023), is an interesting finding and could have implications for unique GAD65^+^-only cell function.

### Implications of bipotential plasticity

VTA GABAergic neurons provide either local interneuron inhibition of VTA DA neurons or long-range projections to diverse brain regions (Bouarab *et al*., 2019). Local VTA GABA interneuron connections make up nearly half of all inputs to VTA DA neurons (Beier, 2022). Functional implications of VTA GABA neurons include mediators of reward and aversion, with distinct behavioral roles for locally-versus distally-derived GABAergic inputs (Bouarab *et al*., 2019; Simon et al., 2024). Indeed, optogenetic activation of VTA GABA interneurons disrupts reward consummatory behavior while leaving cue driven anticipatory behavior unaffected by suppressing VTA DA cell activity (van Zessen *et al*., 2012). New data also illustrates VTA GABA cells encode salience of both rewarding and aversive stimuli (Lefner and Moghaddam, 2025). In addition, VTA GABA cell activity is coordinated in part by regulation via their selective GABAergic inputs that arise from areas including RMTg and LH, with a bias toward inhibitory inputs arriving to the VTA synapsing with local GABA neurons rather than DA cells (Simon *et al*., 2024). Thus, VTA GABA cells are strongly implicated in DA cell function and reward behavior, and their activity is regulated by their inhibitory inputs.

For example, the LH is the feeding center (Anand and Brobeck, 1951), with LH GABAergic neuron activation motivating consumption and appetite, independent of orexin containing LH neurons (Jennings et al., 2015), and their projections to the VTA are key for reward function in measurable behavioral-relevant manners (Nieh et al., 2016). Again, the majority of LH GABAergic input to the VTA innervates VTA GABA interneurons versus DA neurons, leading to disinhibition of DA neurons, and this circuit is often implicated in reward feeding behaviors such as sucrose seeking for reward consumption that goes beyond normal satiety or hunger feeding (Marino et al., 2020; Nieh *et al*., 2016). GABAergic LH inputs to VTA are essential for storing and recalling reward-predictive cues as optogenetic inhibition of LH GABA cells disrupt cue learning and cue-induced food seeking after learning (Sharpe et al., 2017). Indeed, optogenetic activation of LH GABAergic projections to VTA led to enhanced reward behaviors, supporting positive reinforcement (Nieh *et al*., 2016), though GABAergic projections from LH may also target overeating via the locus coeruleus (pons) GABA neurons (Marino *et al*., 2020). Relevant to our study, a 40Hz LH stimulus induced reward while a 5Hz stimulus (similar to our stimulation paradigm) induced GABA-dependent feeding behavior (Barbano et al., 2016), suggesting iLTP induced by our 5 Hz stimulation via the LH pathway could have implications in regulating reward feeding behaviors.

RMTg GABAergic projections mainly innervate DA cells (80%), with a minority of projections innervating non-DAergic cells (Balcita-Pedicino et al., 2011). While RMTg GABA inputs to DA cells can be disinhibited by activation of presynaptic opioid or cannabinoid receptor mechanisms, leading to increased DA cells firing (Lecca et al., 2012; Matsui and Williams, 2011), little to nothing has been reported on the functional implications of RMTg inputs to non-DAergic cells. We confirm the presence of these RMTg inputs to non-dopaminergic and provide evidence that these cells include VTA GABA cells and demonstrate RMTg ability to inhibit and induce iLTP in these cells. Nothing is currently known regarding the implication of a RMTg to VTA GABA cell circuit on reward behavior, though iLTP of this circuit could also theoretically lead to disinhibition of DA neurons leading to reward behaviors. Further studies will need to elucidate their role in reward behavior.

Regarding functional implications of local VTA GABA cell interneurons that are selective for other VTA GABA cells there is little to nothing known. In addition, the impact of cocaine-occlusion of iLTD would depend on the postsynaptic output of VTA GABA cells, which likely innervate DA neurons and thus influence reward-behavior or reward-prediction. If iLTD was induced (i.e. occlusion) by cocaine it would suggest increased VTA GABA cell activation and thus a hypodopaminergic state, while elimination of iLTD capacity would do the opposite.

The former is more likely to occur, due to the withdrawal state cocaine induces and thus could play a role in drug-seeking.

### Cocaine and GABA_B_-dependent iLTD

Both acute and repeated use of cocaine alter synaptic plasticity in various brain areas, including synapses onto VTA DA and GABA neurons (Bocklisch *et al*., 2013; Friend et al., 2021; Ungless *et al*., 2001; Wolf, 2016). In this study, we identified cocaine-induced elimination of iLTD, while iLTP was resistant and spared, suggesting a differential impact of cocaine on plasticity of GABA cells that could have implications on the VTA inhibitory circuit in response to cocaine. As iLTD is mediated by local VTA inhibitory inputs, it suggests that local inputs are more sensitive to cocaine than projecting inputs from LH/RMTg. Selective iLTD elimination suggests greater impact on DA function in reward behavior (Brown *et al*., 2012; Tan *et al*., 2012), than reward feeding from LH or behavioral impact from the RMTg.

Interestingly, a prior study demonstrated high-frequency (50Hz) stimulus-induced iLTP from NAc inputs from D_1_-expressing inhibitory inputs from medium spiny neurons to VTA GABA cells was occluded by repeated administration of cocaine, thereby disinhibiting VTA DA neurons (Bocklisch *et al*., 2013). Our results are seemingly in conflict with this observation; however, we do not know if there are mechanistically unique iLTP types to VTA GABA neurons (i.e. NAc inputs versus LH or RMTg inputs), and there may be as we noted NMDA-receptor dependence of iLTP in some GABA cells, but not others. In addition, our results indicate iLTP from RMTg inputs was significantly larger than LH inputs and note iLTP from NAc inputs was sensitive to cocaine exposure (Bocklisch *et al*., 2013). Therefore, there are possibly two or more iLTP mechanisms (based on inputs or conditioning stimulus employed), one sensitive to cocaine, and the others not, with differential NMDA receptor sensitivity.

We previously determined iLTD was GABA_B_ receptor-dependent (Nufer *et al*., 2023). Cocaine occlusion of iLTD here is consistent with other reports of both acute and chronic treatments of cocaine/psychostimulants suppression of transmission via GABA_B_ receptors (Arora et al., 2011; Padgett et al., 2012; Sharpe et al., 2014). These suggest one possibility of reduced GABA_B_ impact on neurotransmission is due to the functional loss of G-Protein-coupled inwardly rectifying K^+^ (GIRK) conductance, likely via an intracellular Ca^2+^ mechanism (Sharpe *et al*., 2014), or alternatively to reduce GABA_B_R synaptic currents by increased adenosine tone (Bonci and Williams, 1996). The restoration of chronic cocaine-eliminated iLTD by A1 adenosine receptor antagonist we see here suggests a similar mechanism to the latter. Our data also demonstrates the ubiquitous presence of A1 at GABAergic inputs to VTA GABA cells as GABAergic input was depressed in all cells. As the activation of A1 decreases cAMP, and resveratrol, a phytoalexin that elevates cAMP also attenuates cocaine-induced conditioned place preference (Li et al., 2017), this is a promising target for cocaine dependence treatment. Lastly, as both alcohol (Nufer *et al*., 2023) and cocaine block iLTD with optogenetic data demonstrating iLTD is mediated by local VTA GABA inputs, it suggests VTA GABA cells are those impacted and dysfunctional following cocaine or alcohol exposure, compared to GABA inputs projecting to the VTA (with the exception of NAc (Bocklisch *et al*., 2013). This suggests local VTA GABA cells synaptic connections as those needing further examination for potential dependent-treatment strategies.

## Conclusion

This research builds upon our prior findings of mechanisms governing inhibitory VTA reward circuit plasticity and the impact of psychostimulant cocaine on this plasticity. Here we note for the first time that bipotential plasticity to an inhibitory cell is influenced by their regional inputs. Since cocaine can affect plasticity in multiple brain regions, improving the odds of treating cocaine addiction lies in a greater understanding of cocaine impact on the entire reward circuit. Our exploration of the intricate GABA circuit upstream of the VTA DA neurons will add a crucial piece to the puzzle of how cocaine modifies the reward system. By integrating our insights with ongoing clinical trials and pharmacological research, we will move closer to treatments that will reverse cocaine dependence, which is proving elusive (Czoty et al., 2016).

## Acknowledgements

NIH National Institute of Drug Abuse grants R15DA038092 (JE) and R15DA049260 (JE) funded this work. The content is solely the responsibility of the authors and does not necessarily represent the official views of the National Institutes of Health. This work was also supported by institutional Mentoring Grants (JE). We thank the NIDA Drug Supply Program for providing the cocaine used in this study. We also acknowledge assistance of Annie Warner, Ben Daynes, and Stockton Porter for assistance with IHC, as well as Scott Steffensen for providing GAD65-mCherry mice. We also thank Dr. Karl Deisseroth and UNC Vector Core for providing the AAV-associated ChR2-mCherry virus for optogenetic experiments.

## Author Contributions

BJW and JGE designed the research and authored the paper. BJW and SH performed electrophysiology experiments. HE, IF, DB, and CB performed immunohistochemistry experiments and helped edit the paper. JE provided funding and resources for the project. All authors performed the studies and analyzed the data, edited the paper, and approve the final version.

## Reference

1. Anand, B.K., and Brobeck, J.R. (1951). Localization of a “feeding center” in the hypothalamus of the rat. Proc Soc Exp Biol Med 77, 323–324. 10.3181/00379727-77-18766.

2. Arora, D., Hearing, M., Haluk, D.M., Mirkovic, K., Fajardo-Serrano, A., Wessendorf, M.W., Watanabe, M., Luján, R., and Wickman, K. (2011). Acute cocaine exposure weakens GABA(B) receptor-dependent G-protein-gated inwardly rectifying K+ signaling in dopamine neurons of the ventral tegmental area. The Journal of neuroscience: the official journal of the Society for Neuroscience 31, 12251–12257. 10.1523/jneurosci.0494-11.2011.

3. Balcita-Pedicino, J.J., Omelchenko, N., Bell, R., and Sesack, S.R. (2011). The inhibitory influence of the lateral habenula on midbrain dopamine cells: ultrastructural evidence for indirect mediation via the rostromedial mesopontine tegmental nucleus. The Journal of comparative neurology 519, 1143–1164. 10.1002/cne.22561.

4. Barbano, M.F., Wang, H.L., Morales, M., and Wise, R.A. (2016). Feeding and Reward Are Differentially Induced by Activating GABAergic Lateral Hypothalamic Projections to VTA. The Journal of neuroscience: the official journal of the Society for Neuroscience 36, 2975–2985. 10.1523/jneurosci.3799-15.2016.

5. Barker, D.J., Root, D.H., Zhang, S., and Morales, M. (2016). Multiplexed neurochemical signaling by neurons of the ventral tegmental area. Journal of chemical neuroanatomy 73, 33–42. 10.1016/j.jchemneu.2015.12.016.

6. Beier, K. (2022). Modified viral-genetic mapping reveals local and global connectivity relationships of ventral tegmental area dopamine cells. Elife 11. 10.7554/eLife.76886.

7. Bocklisch, C., Pascoli, V., Wong, J.C., House, D.R., Yvon, C., de Roo, M., Tan, K.R., and Lüscher, C. (2013). Cocaine disinhibits dopamine neurons by potentiation of GABA transmission in the ventral tegmental area. Science 341, 1521–1525. 10.1126/science.1237059.

8. Bonci, A., and Williams, J.T. (1996). A common mechanism mediates long-term changes in synaptic transmission after chronic cocaine and morphine. Neuron 16, 631–639. 10.1016/s0896-6273(00)80082-3.

9. Bouarab, C., Thompson, B., and Polter, A.M. (2019). VTA GABA Neurons at the Interface of Stress and Reward. Frontiers in neural circuits 13. 10.3389/fncir.2019.00078.

10. Brischoux, F., Chakraborty, S., Brierley, D.I., and Ungless, M.A. (2009). Phasic excitation of dopamine neurons in ventral VTA by noxious stimuli. Proceedings of the National Academy of Sciences of the United States of America 106, 4894–4899. 10.1073/pnas.0811507106.

11. Brown, M.T.C., Tan, K.R., O’Connor, E.C., Nikonenko, I., Muller, D., and Luscher, C. (2012). Ventral tegmental area GABA projections pause accumbal cholinergic interneurons to enhance associative learning. Nature 492, 452–456. http://www.nature.com/nature/journal/v492/n7429/abs/nature11657.html#supplementary-information.

12. Chen, X., Li, X., Wong, Y.T., Zheng, X., Wang, H., Peng, Y., Feng, H., Feng, J., Baibado, J.T., Jesky, R., et al. (2019). Cholecystokinin release triggered by NMDA receptors produces LTP and sound-sound associative memory. Proceedings of the National Academy of Sciences of the United States of America 116, 6397–6406. 10.1073/pnas.1816833116.

13. Chieng, B., Azriel, Y., Mohammadi, S., and Christie, M.J. (2011). Distinct cellular properties of identified dopaminergic and GABAergic neurons in the mouse ventral tegmental area. Journal of Physiology-London 589, 3775–3787. 10.1113/jphysiol.2011.210807.

14. Crawley, J.N. (1985). Comparative distribution of cholecystokinin and other neuropeptides. Why is this peptide different from all other peptides? Annals of the New York Academy of Sciences 448, 1–8. 10.1111/j.1749-6632.1985.tb29900.x.

15. Creed, M.C., Ntamati, N.R., and Tan, K.R. (2014). VTA GABA neurons modulate specific learning behaviors through the control of dopamine and cholinergic systems. Frontiers in behavioral neuroscience 8, 8. 10.3389/fnbeh.2014.00008.

16. Czoty, P.W., Stoops, W.W., and Rush, C.R. (2016). Evaluation of the “Pipeline” for Development of Medications for Cocaine Use Disorder: A Review of Translational Preclinical, Human Laboratory, and Clinical Trial Research. Pharmacological reviews 68, 533–562. 10.1124/pr.115.011668.

17. Dacher, M., and Nugent, F.S. (2011). Morphine-induced modulation of LTD at GABAergic synapses in the ventral tegmental area. Neuropharmacology 61, 1166–1171. 10.1016/j.neuropharm.2010.11.012.

18. Di Ciano, P., and Everitt, B.J. (2003). The GABA(B) receptor agonist baclofen attenuates cocaine– and heroin-seeking behavior by rats. Neuropsychopharmacology 28, 510–518. 10.1038/sj.npp.1300088.

19. Dobi, A., Margolis, E.B., Wang, H.L., Harvey, B.K., and Morales, M. (2010). Glutamatergic and nonglutamatergic neurons of the ventral tegmental area establish local synaptic contacts with dopaminergic and nondopaminergic neurons. The Journal of neuroscience: the official journal of the Society for Neuroscience 30, 218–229. 10.1523/jneurosci.3884-09.2010.

20. Edwards, N.J., Tejeda, H.A., Pignatelli, M., Zhang, S., McDevitt, R.A., Wu, J., Bass, C.E., Bettler, B., Morales, M., and Bonci, A. (2017). Circuit specificity in the inhibitory architecture of the VTA regulates cocaine-induced behavior. Nat Neurosci 20, 438–448. 10.1038/nn.4482.

21. Erhardt, S., Mathé, J.M., Chergui, K., Engberg, G., and Svensson, T.H. (2002). GABA(B) receptor-mediated modulation of the firing pattern of ventral tegmental area dopamine neurons in vivo. Naunyn-Schmiedeberg’s archives of pharmacology 365, 173–180. 10.1007/s00210-001-0519-5.

22. Faget, L., Osakada, F., Duan, J., Ressler, R., Johnson, Alexander B., Proudfoot, James A., Yoo, Ji H., Callaway, Edward M., and Hnasko, Thomas S. (2016). Afferent Inputs to Neurotransmitter-Defined Cell Types in the Ventral Tegmental Area. Cell Reports 15, 2796–2808. 10.1016/j.celrep.2016.05.057.

23. Friend, L.N., Wu, B., and Edwards, J.G. (2021). Acute cocaine exposure occludes long-term depression in ventral tegmental area GABA neurons. Neurochem Int, 105002. 10.1016/j.neuint.2021.105002.

24. Gonzalez-Hernandez, T., Barroso-Chinea, P., Acevedo, A., Salido, E., and Rodriguez, M. (2001). Colocalization of tyrosine hydroxylase and GAD65 mRNA in mesostriatal neurons. The European journal of neuroscience 13, 57–67.

25. Jayaram, P., and Steketee, J.D. (2004). Effects of repeated cocaine on medial prefrontal cortical GABAB receptor modulation of neurotransmission in the mesocorticolimbic dopamine system. Journal of neurochemistry 90, 839–847. 10.1111/j.1471-4159.2004.02525.x.

26. Jennings, J.H., Ung, R.L., Resendez, S.L., Stamatakis, A.M., Taylor, J.G., Huang, J., Veleta, K., Kantak, P.A., Aita, M., Shilling-Scrivo, K., et al. (2015). Visualizing hypothalamic network dynamics for appetitive and consummatory behaviors. Cell 160, 516–527. 10.1016/j.cell.2014.12.026.

27. Jhou, T.C., Geisler, S., Marinelli, M., Degarmo, B.A., and Zahm, D.S. (2009). The mesopontine rostromedial tegmental nucleus: A structure targeted by the lateral habenula that projects to the ventral tegmental area of Tsai and substantia nigra compacta. The Journal of comparative neurology 513, 566–596. 10.1002/cne.21891.

28. Juarez, B., and Han, M.H. (2016). Diversity of Dopaminergic Neural Circuits in Response to Drug Exposure. Neuropsychopharmacology 41, 2424–2446. 10.1038/npp.2016.32.

29. Kasanetz, F., Deroche-Gamonet, V., Berson, N., Balado, E., Lafourcade, M., Manzoni, O., and Piazza, P.V. (2010). Transition to addiction is associated with a persistent impairment in synaptic plasticity. Science 328, 1709–1712. 10.1126/science.1187801.

30. Kaufling, J., Veinante, P., Pawlowski, S.A., Freund-Mercier, M.J., and Barrot, M. (2010). gamma-Aminobutyric acid cells with cocaine-induced DeltaFosB in the ventral tegmental area innervate mesolimbic neurons. Biol Psychiatry 67, 88–92. 10.1016/j.biopsych.2009.08.001.

31. Klitenick, M.A., DeWitte, P., and Kalivas, P.W. (1992). Regulation of somatodendritic dopamine release in the ventral tegmental area by opioids and GABA: an in vivo microdialysis study. The Journal of neuroscience: the official journal of the Society for Neuroscience 12, 2623–2632. 10.1523/jneurosci.12-07-02623.1992.

32. Koob, G.F., and Volkow, N.D. (2010). Neurocircuitry of addiction. Neuropsychopharmacology 35, 217–238. 10.1038/npp.2009.110.

33. Lassen, M.B., Brown, J.E., Stobbs, S.H., Gunderson, S.H., Maes, L., Valenzuela, C.F., Ray, A.P., Henriksen, S.J., and Steffensen, S.C. (2007). Brain stimulation reward is integrated by a network of electrically coupled GABA neurons. Brain research 1156, 46–58. 10.1016/j.brainres.2007.04.053.

34. Lecca, S., Melis, M., Luchicchi, A., Muntoni, A.L., and Pistis, M. (2012). Inhibitory inputs from rostromedial tegmental neurons regulate spontaneous activity of midbrain dopamine cells and their responses to drugs of abuse. Neuropsychopharmacology 37, 1164–1176. 10.1038/npp.2011.302.

35. Lefner, M.J., and Moghaddam, B. (2025). Flexible updating of reward and punishment contingencies by VTA GABA neurons. Curr Biol. 10.1016/j.cub.2025.07.021.

36. Li, Y., Yu, L., Zhao, L., Zeng, F., and Liu, Q.S. (2017). Resveratrol modulates cocaine-induced inhibitory synaptic plasticity in VTA dopamine neurons by inhibiting phosphodiesterases (PDEs). Scientific reports 7, 15657. 10.1038/s41598-017-16034-9.

37. Luscher, C., and Bellone, C. (2008). Cocaine-evoked synaptic plasticity: a key to addiction? Nat Neurosci 11, 737–738. 10.1038/nn0708-737.

38. Lüscher, C., and Malenka, R.C. (2011). Drug-evoked synaptic plasticity in addiction: from molecular changes to circuit remodeling. Neuron 69, 650–663. 10.1016/j.neuron.2011.01.017.

39. Lüscher, C., and Ungless, M.A. (2006). The mechanistic classification of addictive drugs. PLoS Med 3, e437. 10.1371/journal.pmed.0030437.

40. Margolis, E.B., Toy, B., Himmels, P., Morales, M., and Fields, H.L. (2012). Identification of Rat Ventral Tegmental Area GABAergic Neurons. PloS one 7, e42365. 10.1371/journal.pone.0042365.

41. Marino, R.A.M., McDevitt, R.A., Gantz, S.C., Shen, H., Pignatelli, M., Xin, W., Wise, R.A., and Bonci, A. (2020). Control of food approach and eating by a GABAergic projection from lateral hypothalamus to dorsal pons. Proceedings of the National Academy of Sciences of the United States of America 117, 8611–8615. 10.1073/pnas.1909340117.

42. Martinez Damonte, V., Pomrenze, M.B., Manning, C.E., Casper, C., Wolfden, A.L., Malenka, R.C., and Kauer, J.A. (2023). Somatodendritic Release of Cholecystokinin Potentiates GABAergic Synapses Onto Ventral Tegmental Area Dopamine Cells. Biol Psychiatry 93, 197–208. 10.1016/j.biopsych.2022.06.011.

43. Matsui, A., and Williams, J.T. (2011). Opioid-sensitive GABA inputs from rostromedial tegmental nucleus synapse onto midbrain dopamine neurons. The Journal of neuroscience: the official journal of the Society for Neuroscience 31, 17729–17735. 10.1523/jneurosci.4570-11.2011.

44. Merrill, C.B., Friend, L.N., Newton, S.T., Hopkins, Z.H., and Edwards, J.G. (2015). Ventral tegmental area dopamine and GABA neurons: Physiological properties and expression of mRNA for endocannabinoid biosynthetic elements. Scientific reports 5, 16176. 10.1038/srep16176.

45. Morales, M., and Margolis, E.B. (2017). Ventral tegmental area: cellular heterogeneity, connectivity and behaviour. Nature reviews. Neuroscience 18, 73–85. 10.1038/nrn.2016.165.

46. Nieh, E.H., Vander Weele, C.M., Matthews, G.A., Presbrey, K.N., Wichmann, R., Leppla, C.A., Izadmehr, E.M., and Tye, K.M. (2016). Inhibitory Input from the Lateral Hypothalamus to the Ventral Tegmental Area Disinhibits Dopamine Neurons and Promotes Behavioral Activation. Neuron 90, 1286–1298. 10.1016/j.neuron.2016.04.035.

47. Nufer, T., Wu, B., Boyce, Z., Steffens, S., and Edwards, J.G. (2023). Ethanol Blocks a Novel Form of iLTD, but not iLTP of Inhibitory Inputs to VTA GABA Neurons. Neuropsychopharmacology *In Press*.

48. Nugent, F.S., Penick, E.C., and Kauer, J.A. (2007). Opioids block long-term potentiation of inhibitory synapses. Nature 446, 1086–1090.

49. Olson, V.G., and Nestler, E.J. (2007). Topographical organization of GABAergic neurons within the ventral tegmental area of the rat. Synapse (New York, N.Y.) 61, 87–95. 10.1002/syn.20345.

50. Omelchenko, N., Bell, R., and Sesack, S.R. (2009). Lateral habenula projections to dopamine and GABA neurons in the rat ventral tegmental area. European Journal of Neuroscience 30, 1239–1250. 10.1111/j.1460-9568.2009.06924.x.

51. Omelchenko, N., and Sesack, S.R. (2009). Ultrastructural analysis of local collaterals of rat ventral tegmental area neurons: GABA phenotype and synapses onto dopamine and GABA cells. Synapse (New York, N.Y.) 63, 895–906. 10.1002/syn.20668.

52. Padgett, Claire L., Lalive, Arnaud L., Tan, Kelly R., Terunuma, M., Munoz, Michaelanne B., Pangalos, Menelas N., Martínez-Hernández, J., Watanabe, M., Moss, Stephen J., Luján, R., et al. (2012). Methamphetamine-Evoked Depression of GABAB Receptor Signaling in GABA Neurons of the VTA. Neuron 73, 978–989. 10.1016/j.neuron.2011.12.031.

53. Paul, E.J., Tossell, K., and Ungless, M.A. (2019). Transcriptional profiling aligned with in situ expression image analysis reveals mosaically expressed molecular markers for GABA neuron sub-groups in the ventral tegmental area. The European journal of neuroscience 50, 3732–3749. 10.1111/ejn.14534.

54. Phillips, R.A., 3rd, Tuscher, J.J., Black, S.L., Andraka, E., Fitzgerald, N.D., Ianov, L., and Day, J.J. (2022). An atlas of transcriptionally defined cell populations in the rat ventral tegmental area. Cell Rep 39, 110616. 10.1016/j.celrep.2022.110616.

55. Rehfeld, J.F. (2017). Cholecystokinin-From Local Gut Hormone to Ubiquitous Messenger. Frontiers in endocrinology 8, 47. 10.3389/fendo.2017.00047.

56. Root, D.H., Barker, D.J., Estrin, D.J., Miranda-Barrientos, J.A., Liu, B., Zhang, S., Wang, H.L., Vautier, F., Ramakrishnan, C., Kim, Y.S., et al. (2020). Distinct Signaling by Ventral Tegmental Area Glutamate, GABA, and Combinatorial Glutamate-GABA Neurons in Motivated Behavior. Cell Rep 32, 108094. 10.1016/j.celrep.2020.108094.

57. Sharpe, A.L., Varela, E., Bettinger, L., and Beckstead, M.J. (2014). Methamphetamine self-administration in mice decreases GIRK channel-mediated currents in midbrain dopamine neurons. The international journal of neuropsychopharmacology 18. 10.1093/ijnp/pyu073.

58. Sharpe, M.J., Marchant, N.J., Whitaker, L.R., Richie, C.T., Zhang, Y.J., Campbell, E.J., Koivula, P.P., Necarsulmer, J.C., Mejias-Aponte, C., Morales, M., et al. (2017). Lateral Hypothalamic GABAergic Neurons Encode Reward Predictions that Are Relayed to the Ventral Tegmental Area to Regulate Learning. Curr Biol 27, 2089–2100.e2085. 10.1016/j.cub.2017.06.024.

59. Shen, H., and Kalivas, P.W. (2013). Reduced LTP and LTD in prefrontal cortex synapses in the nucleus accumbens after heroin self-administration. The international journal of neuropsychopharmacology 16, 1165–1167. 10.1017/s1461145712001071.

60. Simon, R.C., Loveless, M.C., Yee, J.X., Goh, B., Cho, S.G., Nasir, Z., Hashikawa, K., Stuber, G.D., Zweifel, L.S., and Soden, M.E. (2024). Opto-seq reveals input-specific immediate-early gene induction in ventral tegmental area cell types. Neuron. 10.1016/j.neuron.2024.05.026.

61. St Laurent, R., and Kauer, J. (2019). Synaptic Plasticity at Inhibitory Synapses in the Ventral Tegmental Area Depends upon Stimulation Site. eNeuro 6. 10.1523/eneuro.0137-19.2019.

62. Stamatakis, Alice M., Jennings, Joshua H., Ung, Randall L., Blair, Grace A., Weinberg, Richard J., Neve, Rachael L., Boyce, F., Mattis, J., Ramakrishnan, C., Deisseroth, K., and Stuber, Garret D. (2013). A Unique Population of Ventral Tegmental Area Neurons Inhibits the Lateral Habenula to Promote Reward. Neuron 80, 1039–1053. 10.1016/j.neuron.2013.08.023.

63. Su, J., Huang, F., Tian, Y., Tian, R., Qianqian, G., Bello, S.T., Zeng, D., Jendrichovsky, P., Lau, C.G., Xiong, W., et al. (2023). Entorhinohippocampal cholecystokinin modulates spatial learning by facilitating neuroplasticity of hippocampal CA3-CA1 synapses. Cell Rep 42, 113467. 10.1016/j.celrep.2023.113467.

64. Tan, Kelly R., Yvon, C., Turiault, M., Mirzabekov, Julie J., Doehner, J., Labouèbe, G., Deisseroth, K., Tye, Kay M., and Lüscher, C. (2012). GABA Neurons of the VTA Drive Conditioned Place Aversion. Neuron 73, 1173–1183. 10.1016/j.neuron.2012.02.015.

65. Ting, J.T., Lee, B.R., Chong, P., Soler-Llavina, G., Cobbs, C., Koch, C., Zeng, H., and Lein, E. (2018). Preparation of Acute Brain Slices Using an Optimized N-Methyl-D-glucamine Protective Recovery Method. J Vis Exp. 10.3791/53825.

66. Ungless, M.A., Whistler, J.L., Malenka, R.C., and Bonci, A. (2001). Single cocaine exposure in vivo induces long-term potentiation in dopamine neurons. Nature 411, 583–587.

67. van Zessen, R., Phillips, J.L., Budygin, E.A., and Stuber, G.D. (2012). Activation of VTA GABA neurons disrupts reward consumption. Neuron 73, 1184–1194. S0896-6273(12)00177-8 [pii] 10.1016/j.neuron.2012.02.016.

68. Wen, D., Zang, G., Sun, D., Yu, F., Mei, D., Ma, C., and Cong, B. (2014). Cholecystokinin-octapeptide restored morphine-induced hippocampal long-term potentiation impairment in rats. Neuroscience letters 559, 76–81. 10.1016/j.neulet.2013.11.043.

69. Wolf, M.E. (2016). Synaptic mechanisms underlying persistent cocaine craving. Nature reviews. Neuroscience 17, 351–365. 10.1038/nrn.2016.39.

70. Yamaguchi, M., Suzuki, T., Abe, S., Baba, A., Hori, T., and Okado, N. (2002). Repeated cocaine administration increases GABA(B(1)) subunit mRNA in rat brain. Synapse (New York, N.Y.) 43, 175–180. 10.1002/syn.10037.

71. Zhou, K., Xu, H., Lu, S., Jiang, S., Hou, G., Deng, X., He, M., and Zhu, Y. (2022). Reward and aversion processing by input-defined parallel nucleus accumbens circuits in mice. Nat Commun 13, 6244. 10.1038/s41467-022-33843-3.

